# Tbx2 is essential for cochlear inner hair cell development and regeneration

**DOI:** 10.1101/2022.05.05.490782

**Authors:** Zhenghong Bi, Xiang Li, Minhui Ren, Yunpeng Gu, Tong Zhu, Shuting Li, Guangqin Wang, Suhong Sun, Yuwei Sun, Zhiyong Liu

**Author notes:** These authors contributed equally to this work (Zhenghong Bi, Xiang Li, Minhui Ren).

## Abstract

Atoh1 is essential for the development of outer hair cells (OHCs) and inner hair cells (IHCs) in the mammalian cochlea. Whereas Ikzf2 is necessary for OHC development, the key gene for IHC development remains unknown. We found that deleting Tbx2 in neonatal IHCs led to their trans-differentiation into OHCs, by repressing 26.7% of IHC and inducing 56.3% of OHC genes, including *Ikzf2*. More importantly, persistent expression of Tbx2 together with transient Atoh1 effectively reprogramed non-sensory supporting cells into new IHCs expressing functional IHC marker vGlut3. The differentiation status of these new IHCs is much more advanced than those previously reported. Thus, Tbx2 is essential for IHC development, and its co-upregulation with Atoh1 in supporting cells represents a new approach for treating IHC degeneration-related deafness.

## Introduction

Precisely specifying different cell fates is crucial for organogenesis. The mouse cochlea has been widely used as a fascinating model to decipher the molecular mechanisms underlying this fundamental event, because it contains two simple sound receptor cells: inner hair cells (IHCs) and outer hair cells (OHCs) ^1–3^. IHCs and OHCs are derived from the same progenitors expressing *Atoh1*, a key b-HLH transcriptional factor (TF) needed in generating both IHCs and OHCs ^4,5^. Adjacent to HCs are different subtypes of non-sensory cochlear supporting cells (SCs), from medial to lateral, which are known as Inner border cells (IBCs), Inner phalangeal cells (IPhs), Pillar cells (PCs) and Deiters’ cells (DCs) ^1,2,6^. The cochlear SCs of non-mammalian vertebrates including birds and fish are able to regenerate HCs upon damage, however, SCs in mammals lose such a regenerative capacity ^7,8^. Therefore, damaging or degeneration of either OHCs or IHCs would result in permanent hearing impairment in mammals including human beings.

The IHCs and OHCs share many pan-HC markers such as *Myo7a,* but they also differ in many aspects. OHCs are sound amplifiers and specifically express motor protein Prestin encoded by *Slc26a5 ^9,10^*, and *Slc26a5^-/-^* mice display severe hearing impairment ^11^. IHCs are primary sensory cells and specifically express vGlut3 encoded by *Slc17a8*, Otoferlin and Slc7a14 ^12–15^. The *Slc17a8^-/-^ and Otoferlin ^-/-^* mice are profoundly deaf because both vGlut3 and Otoferlin are heavily involved in packaging and exocytosis of ribbon synapse vesicles containing the excitatory neurotransmitter glutamate in the IHCs. Recently, it is reported that *Insm1* and *Ikzf2* are the key regulators of OHC development and accordingly OHCs tend to transdifferentiate into IHCs or IHC-like cells in *Insm1* or *Ikzf2* mutant mice^16,17^.

In contrast, it remains poorly understood regarding what gene is needed for normal IHC development as well as what gene, together with *Atoh1*, are capable of regenerating vGlut3+ IHCs from cochlear SCs, especially from the IBCs/IPhs that are close to IHCs and have the intrinsic plasticity to transdifferentiate into IHCs, as reported in our previous study^18^. Addressing this question would undoubtedly provide new insights into the molecular mechanisms underlying IHC development and regeneration. In this study we identified that T-box transcription factor 2 (*Tbx2*) is the transcriptional factor (TF) highly and persistently expressed in IHCs, but not OHCs. When *Tbx2* is conditionally deleted, most if not all IHCs tend to transdifferentiate into OHCs, with upregulating 56.3% of OHC and downregulating 26.7% of IHC genes, respectively. As far as we are aware of, *Tbx2* is the first gene reported to be necessary in IHC development. Besides its critical role in normal IHC development, we demonstrated that, *Tbx2* and *Atoh1* together are sufficient to transform the neonatal cochlear IBCs/IPhs into vGlut3+ new IHCs, and the reprogramming efficiency is ∼29.5%. The new IHCs also express other IHC markers Otoferlin and Slc7a14 and possess the bird wing-like stereocilia. Furthermore, the new IHCs generally follow the developmental pathway of the wild type IHCs, first expressing general HC marker Myo7a, followed by vGlut3. Collectively, our study uncovers the critical roles of Tbx2 in both IHC development and regeneration, and will pave the way for future IHC regeneration.

## Results

### *Tbx2* is highly expressed in IHCs but not in OHCs

We aimed to identify genes that are highly expressed in IHCs, but depleted in OHCs, with hypothesis that they are the key candidate regulators for IHC development. *Slc17a8^iCreER/^*^+^; *Rosa26*-CAG-LSL-Tdtomato (Ai9)/+ (abbreviated as Slc17a8-Ai9) were administered tamoxifen (TMX) at postnatal day 2 (P2) and P3 ^19^. We manually picked 50 Tdtomato+ endogenous IHCs at P30 (P30_WT IHCs) that were subjected to single cell RNA-Seq (scRNA-Seq) via smart-seq approach (Fig. 1A). We compared gene profiles between 50 P30_WT IHCs and 17 wild type OHCs at P30 (P30_WT OHCs) reported in our previous study ^20^, and found 2,162 differentially (FC>2, p<0.05) expressed genes (table S1). With the criteria of transcript per million (TPM) above 16 ^20^, 389 were defined as IHC genes, and 151 were as OHC genes (Fig. 1B and table S1). Those genes included known IHC markers *Slc17a8*, *Tbx2*, *Otof* and *Slc7a14 ^12–15,17^*, as well as OHC markers *Slc26a5*, *Ikzf2*, *Lbh* and *Sri ^10,16,21,22^* (Fig. 1C). Notably, *Tbx2* is the top TF highly expressed in IHCs but not OHCs (red arrow in Fig. 1C). However, Tbx2 protein expression pattern remains poorly characterized in cochleae.

**Fig. 1.**
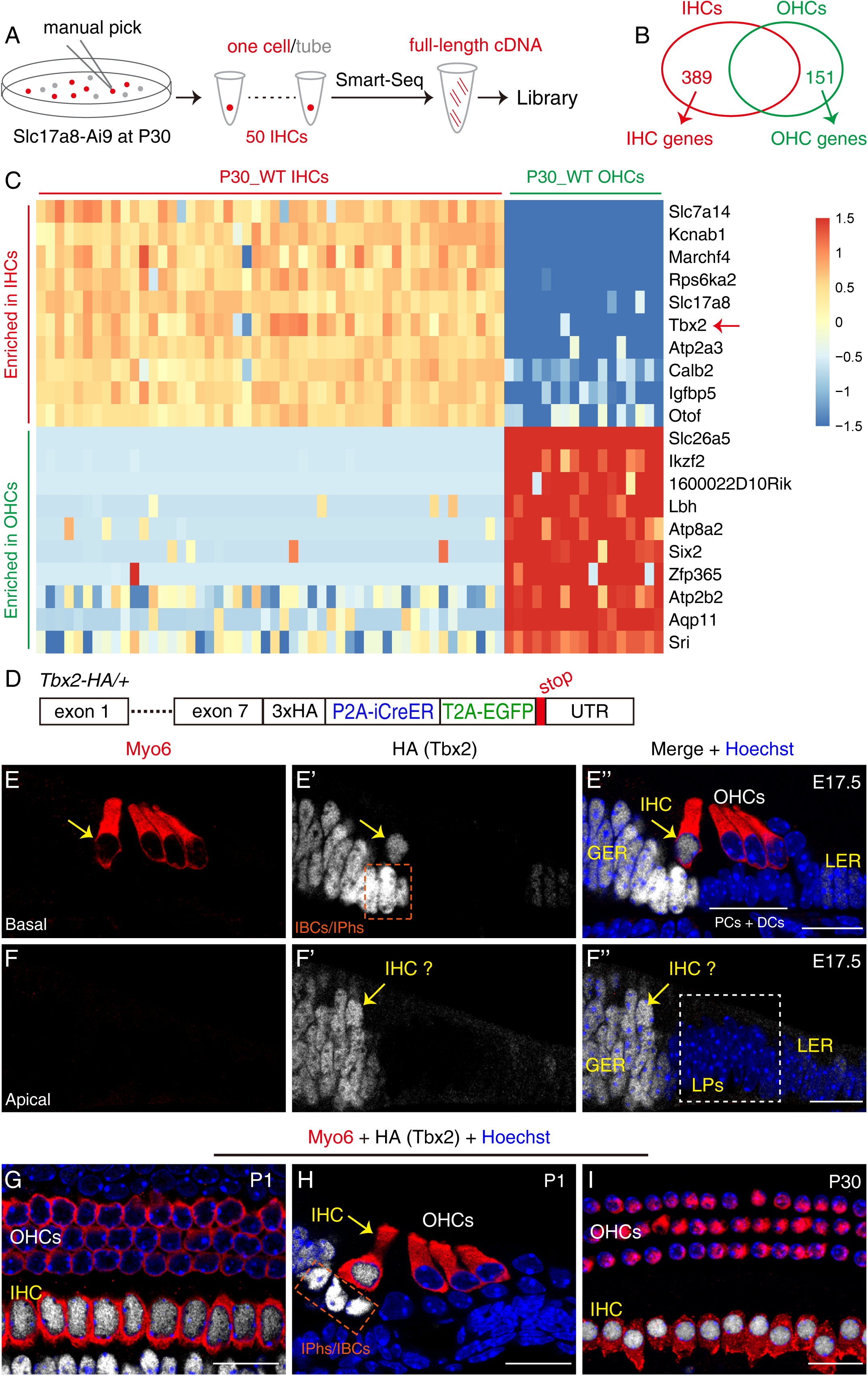
Tbx2 is highly expressed in IHCs, but not in OHCs. **(A-B)** The illustration of single-cell RNA-seq of adult IHCs at P30 via smart-seq (A), 389 IHC and 151 OHC genes are identified (B). **(C)** The heatmap showing the top exampled differently expressed genes between IHCs and OHCs at P30. **(D)** The diagram of *Tbx2*\*3xHA-P2A-iCreER-T2A-EGFP/+ (*Tbx2*-HA/+) and details please refer to fig. S1A-F. **(E-I)** Dual staining of HC marker Myo6 and HA (Tbx2) in cochlear samples of *Tbx2*-HA/+ mice at E17.5 (E-F’’), P1 (G-H) and P30 (I). The yellow arrows in (E-F’’) mark the IHCs. The dotted squares in (E’ and H) label IBCs/IPhs. Scale bar: 20 μm (F’’, G, H, I).

Next, we generated a new knockin mouse strain *Tbx2*\*3xHA-P2A-iCreER-T2A-EGFP/+ (abbreviated as *Tbx2*-HA/+) where three HA tags were fused to the C-terminus of Tbx2 (Fig. 1D and fig. S1A-F). Tbx2 pattern was primarily characterized by HA antibody. In agreement with previous reports ^23,24^, HA (Tbx2) was broadly expressed in otocyst cells expressing Sox2, but not in hindbrain, at embryonic day 9.5 (E9.5) (fig. S1G-G’’). Tbx2 was broadly expressed in cochlear duct cells at E13.5 (fig. S1H). At E15.5, while Tbx2 was maintained in medial portion including Myo6+ IHCs, it became undetected in lateral progenitors (LPs) that would become OHCs, PCs and DCs, in the basal turn (fig. S1I-I’’). Notably, Tbx2 was still broadly expressed in apical turns by E15.5 (fig. S1J-J’’). The wave of Tbx2 downregulation was largely completed by E17.5 when Tbx2 disappeared in basal OHCs, PCs and DCs, and apical LPs (Fig. 1E-F’’). As expected, Tbx2 was highly expressed in IHCs, but not OHCs, at P1 (Fig. 1G and H), P15 and P30 (Fig. 1I). Collectively, Tbx2 is persistently and specifically expressed in both differentiating and mature IHCs, promoting us to hypothesize that Tbx2 is essential for IHC development.

### IHCs transdifferentiate into OHCs when *Tbx2* is conditionally deleted

Our hypothesis was supported by two pieces of genetic evidence. First, we generated a germ line *Tbx2* knockout mouse strain (*Tbx2^+/-^*) in which the entire *Tbx2* loci (∼9.6k bp) was deleted (fig. S2A-C). *Tbx2 ^-/-^* died at early embryonic ages due to cardiac defect ^25^. Notably, OHC specific marker Prestin was weakly detected in *Tbx2^+/-^* but not in *Tbx2^+/+^* IHCs, at P7, P14 and P42. Because the IHC phenotypes in *Tbx2^+/-^* were mild, detailed analysis were not presented, instead, we further generated a *Tbx2 ^flox/+^* mouse strain where the second exon of *Tbx2* was flanked by two loxp sequences (fig. S2D-I). The wild type (WT, control group) and *Slc17a8^iCreER/^*^+^; *Tbx2^flox/-^* (abbreviated as Tbx2 cko, experimental group) mice were administered with tamoxifen (TMX) at P2 and P3.

First, the WT IHCs were vGlut3+/Prestin- (Fig. 2A-A’’), however, ectopic Prestin with heterogeneous levels was detected in IHCs in Tbx2 cko mice (arrows in Fig. 2B-B’’) at P7. Notably, relative to the IHCs without ectopic Prestin (asterisks in Fig. 2B-B’’), vGlut3 expression in Prestin+ IHCs was decreased at different degrees. Secondly, Prestin remained specific in WT OHCs at P14 (Fig. 2C-C’’), but its level in Tbx2cko IHCs was further increased and became comparable to that in the endogenous OHCs (Fig. 2D-D’’). Conversely, vGlut3 expression was still heterogeneous, either was at low levels (blue arrows in Fig. 2D-D’’) or became undetectable (yellow arrows in Fig. 2D-D’’). Moreover, Prestin level was homogenous in majority of the Prestin+ IHCs. It suggested that Prestin increment happened faster than decrement of vGlut3 by P14. As expected, vGlut3 level in IHCs without Prestin (asterisks in Fig. 2D-D’’), was as high as that in the WT IHCs (Fig. 2C-C’’). Thirdly, at P42, relative to WT IHCs (Fig. 2E-E’’), vGlut3 became undetectable in all Prestin+ IHCs (yellow arrows in Fig. 2F-F’’) or was intact in IHCs without Prestin (asterisks in Fig. 2F-F’’). Thus, our data supported that IHCs tended to transdifferentiate into OHCs when *Tbx2* is absent. Hereafter, to differ with the IHCs that did not undergo cell fate change (asterisks in Fig. 2), the Prestin+ IHCs were defined as induced-OHCs derived from IHCs (abbreviated as iOHCs) in which vGlut3 was either decreased vGlut3 or completely lost.

**Fig. 2.**
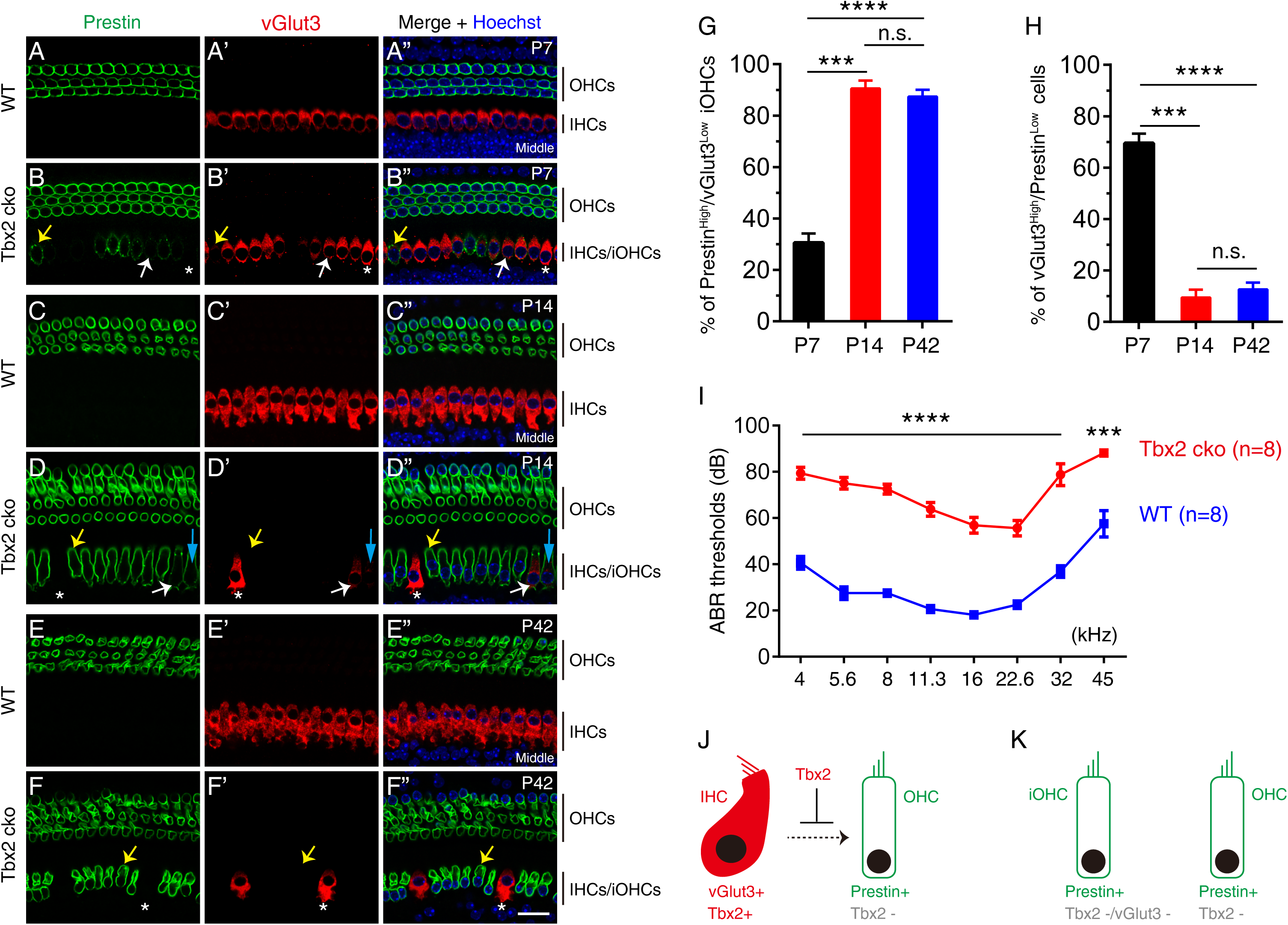
Loss of Tbx2 causes IHCs to gradually decrease vGlut3 but increase Prestin. **(A-F’’)** Dual whole mount staining of vGlut3 and Prestin in cochlear samples from wild type (WT, A-A’’, C-C’’ and E-E’’) and *Tbx2* conditional knockout model (Tbx2 cko, B-B’’, D-D’’ and F-F’’) at P7, P14 and P42, respectively. Yellow arrows mark one Prestin^High^/vGlut3^Low^ iOHC, whereas white arrows label one vGlut3^High^/Prestin^Low^ iOHC. The asterisks represent the IHC that does not undergo cell fate conversion and, to simplify the analysis, is included in the vGlut3^High^/Prestin^Low^ population. **(G-H)** Quantification of the Prestin^High^/vGlut3^Low^ iOHCs (G) and vGlut3^High^/Prestin^Low^ cells (H) in Tbx2 cko mice at P7, P14 and P42. Data are presented as Mean ± SEM. *** p<0.001, **** p<0.0001. n.s.: no significant difference. **(I)** ABR measurements of between wild type (WT, blue line) and Tbx2 cko (red line) mice at P42. Significant difference is detected in all frequencies. **** p<0.0001, *** p<0.001. **(J-K)** The simplified model cartoon to highlight roles of Tbx2 in IHC fate stabilization. After Tbx2 is deleted, IHCs become iOHCs (K). Scale bar: 20 μm.

### The cell fate conversion of iOHCs is largely completed by P14

Next, we estimated the progression of cell fate conversion occurring in those iOHCs between P7 and P42. First, we quantified cells in the entire cochlear turns that expressed high level of Prestin (Prestin^High^), but low (including not detectable) vGlut3 (vGlut3^Low^). Those cells were defined as Prestin^High^/vGlut3^Low^ iOHCs (yellow arrows in Fig. 2B-B’’, yellow and blue arrows in Fig. 2D-D’’ and yellow arrows in Fig. 2F-F’’). The percentages of Prestin^High^/vGlut3^Low^ iOHCs was the lowest (30.5% ± 3.7%) at P7 (n=3), and significantly increased to 90.5% ± 3.1% at P14 (n=3), and 87.4 % ± 2.7 % at P42 (n=3) (Fig. 2G). Although Tbx2 antibody was not available to validate absence of Tbx2, the Prestin^High^/vGlut3^Low^ iOHCs were derived from the endogenous IHCs in which *Tbx2* was successfully deleted. Secondly, we also quantified cells that expressed high level of vGlut3 (vGlut3^High^), but low (also including not detectable) Prestin (Prestin^Low^). Those cells were conversely defined as vGlut3^High^/Prestin^Low^ cells. The percentages of the vGlut3^High^/Prestin^Low^ cells was the highest (69.5% ± 3.7%) at P7, and drastically decreased to 9.5% ± 3.1% at P14, and 12.6 % ± 2.7 % at P42 (Fig. 2H).

Notably, we speculated that vGlut3^High^/Prestin^Low^ cells should include two kinds of subpopulations: 1) the iOHCs that were in the early process of cell fate conversion (white arrows in Fig. 2B-B’’ and Fig. 2D-D’’); 2) the endogenous IHCs that did not undergo cell fate change (asterisks in Fig. 2), either due to their irresponsive to Tbx2 deletion or that Tbx2 was not successfully deleted. Thus, our results suggested that, although the cell fate conversion velocity was not synchronized in different iOHCs, it was pretty completed at P14, because there was no significant difference between P14 and P42 (Fig. 2G and H). Because vGlut3^High^/Prestin^Low^ cells only occupied ∼12.6 % in the IHC region at P42 (Fig. 2H), the hearing thresholds of *Tbx2* cko (n=8) at all frequency regions were significantly (**** p<0.0001 and *** p<0.001) higher than those in WT mice (n=8) (Fig. 2I).

Likewise, we additionally performed double staining of Prestin with two more IHC markers, Otoferlin (fig. S3A-B’’) or Slc7a14 (fig. S3C-D’’), at P42. Briefly, Otoferlin and Slc7a14 exhibited similar dynamics as vGlut3. Moreover, relative to WT IHCs (white circles in fig. S3G-G’’), numbers of ribbon synapses were reduced in the iOHCs. Collectively, we proposed a working model that Tbx2 is essential for IHC development by repressing expression of OHC genes or preventing transdifferentiation of IHCs into OHCs (Fig. 2J). Thus, when the neonatal IHCs lost Tbx2, they gradually became iOHCs (Fig. 2K). The overall degree of cell fate change would be calculated by single-cell transcriptomic analysis.

### The IHC genes are globally repressed and conversely OHC genes are derepressed in the iOHCs

We next thoroughly characterized the transcriptomic profiles of 55 WT IHCs at P14 (P14_WT IHCs) from Slc17a8-Ai9 mice, and the 46 iOHCs at P14 (P14_iOHCs) from *Slc17a8^iCreER/^*^+^; *Tbx2^flox/-^;* Ai9/+ (abbreviated as Slc17a8-Tbx2cko-Ai9) (Fig. 3A and B). Relative to P14_WT IHCs, 862 and 442 genes were significantly (p<0.05, FC>2) upregulated and downregulated, respectively, in P14_iOHCs (Fig. 3C and fig. S4A). All the differentially expressed genes were included in table 2. We noticed that 56.3% (85/151) of the OHC genes were dramatically increased, and that 26.7% (104/389) of the IHC genes were significantly decreased in the P14_iOHCs (Fig. 3D and E). Notably, the upregulated OHC genes included *Slc26a5*, *Lbh* and *Ikzf2* (red arrows in Fig. 3C), and the downregulated IHC genes covered *Otof*, *Slc17a8* and *Slc7a14* (green arrows in Fig. 3C). Collectively, the overall IHC to OHC cell fate conversion degree in the P14_iOHCs was 26.7%-56.3%.

**Fig. 3.**
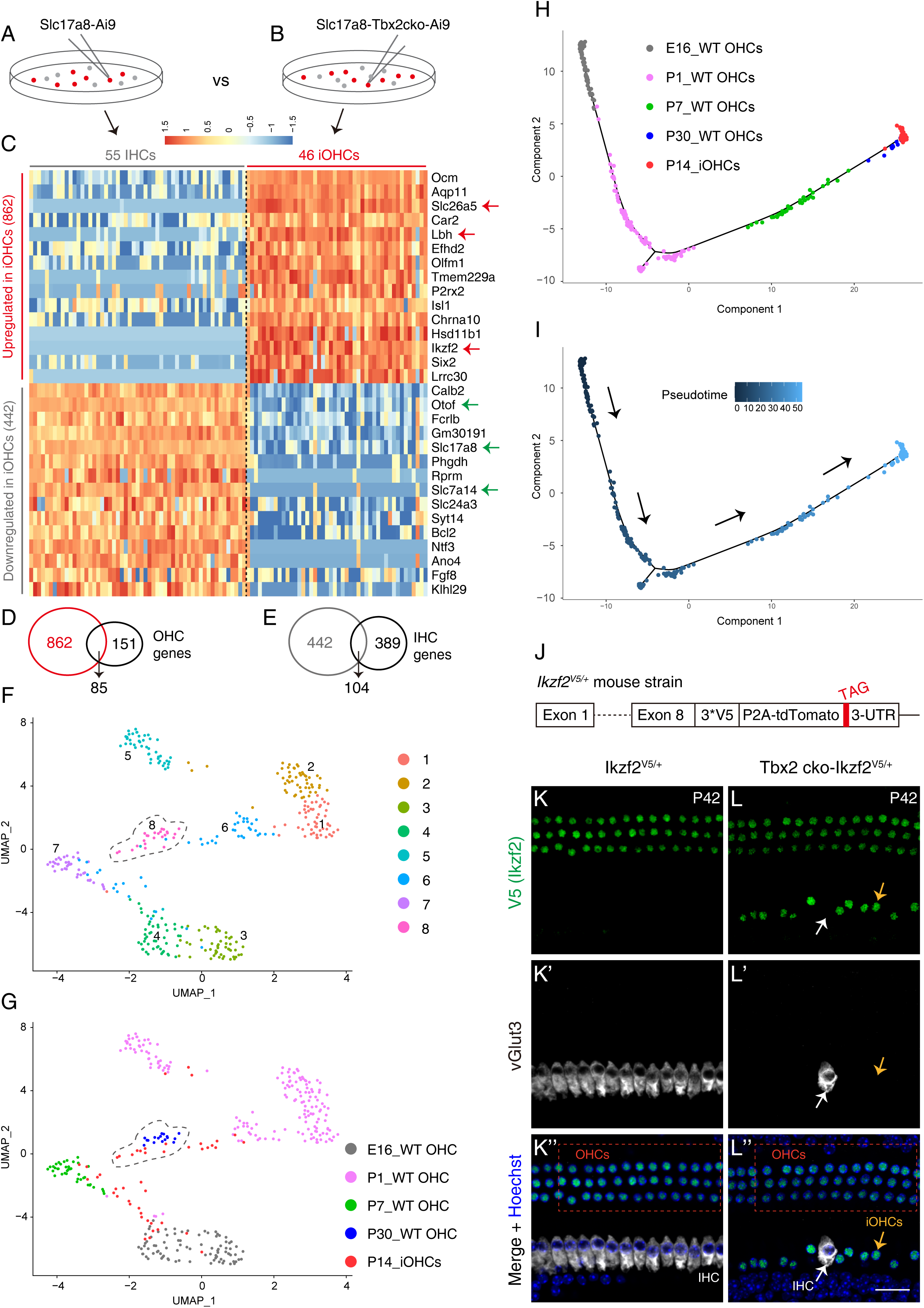
The single-cell transcriptomic profiling of the iOHCs. **(A-B)** Manual picking P14_WT IHCs from Slc17a8-Ai9 mice (A) and P14_iOHCs (B) from Slc17a8-Tbx2cko-Ai9 mice (B). **(C)** Transcriptomic comparison between the P14_WT IHCs and P14_iOHCs. Relative to P14_WT IHCs, 862 and 442genes are significantly upregulated and downregulated in the P14_iOHCs. **(D)** 85 genes including *Slc26a5, Lbh* and *Ikzf2* (red arrows in C) are overlapped between the 862 upregulated and the 151 defined OHC genes. **(E)** 104 genes including *Otof, Slc17a8* and *Slc7a14* (green arrows in C) are overlapped between the 442 downregulated and 389 IHC genes. **(F-G)** UMAP analysis of cell mixture covering E16_WT OHCs, P1_WT OHCs, P7_WT OHCs, P30_WT OHCs, and the P14_iOHCs. 8 main cell clusters are revealed (F). Notably, 7/46 (15.2%) of the P14_iOHCs are assigned to the cluster 8 (gray dotted circle) which primarily comprises of P30_WT OHCs. **(H-I)** Trajectory analysis the same cell mixtures in (F and G). Arrows in (I) represent the calculated developmental ages. Scale bar: 20 μm.

Next, we determined the differentiation status of the endogenous WT OHCs to which the P14_iOHCs were most similar. The 46 P14_iOHCs were pooled with the 87 WT OHCs at E16 (E16_WT OHCs), 170 WT OHCs at P1 (P1_WT OHCs) and 39 WT OHCs at P7 (P7_WT OHCs) from one previous sc-RNA study ^26^, and the 17 P30_WT OHCs ^20^. In total, 8 clusters were revealed by uniform manifold approximation and projection (UMAP) analysis (Fig. 3F). Notably, 7/46 (15.2%) of the P14_iOHCs belonged to the cluster 8 that mainly included the P30_WT OHCs, whereas the remaining 84.8% were assigned to other clusters (Fig. 3G). Moreover, trajectory analysis via monocle demonstrated that P14_iOHCs aggregated with the P30_WT OHCs (Fig. 3H and I), supporting that P14_iOHCs were more similar to P30_WT OHCs than OHCs at other ages. Similar pattern was obtained when additional IHCs, which were obtained from the same sc-RNA study, were included (fig. S4B and C). Together, upon loss of Tbx2, the IHCs became iOHCs that globally resembled, albeit not fully equivalent to, the P30_WT OHCs.

### Ikzf2 protein is also expressed in the iOHCs

Due to the similarity between iOHCs and the endogenous OHCs, we predicted that Ikzf2 protein (also known as Helios), which is a key regulator for OHC development ^16,20^, would be expressed in these iOHCs. Due to lack of good commercial Ikzf2 antibody for immunostaining, we constructed a new knockin mouse strain, *Ikzf2*\*3xV5-P2A-Tdtomato/+ (abbreviated as *Ikzf2^V5/+^*) where three V5 tags were fused with Ikzf2 at its C-terminus (Fig. 3J and fig. S5A-C). The correct gene targeting of *Ikzf2^V5/+^* was confirmed by Southern blot and tail PCR (fig. S5D and E). Thus, V5 antibody could be used to detect Ikzf2 protein. We did not use Tdtomato to represent *Ikzf2* mRNA because its signal to noise ratio was less than V5 antibody.

Co-staining of V5 (Ikzf2) and vGlut3 showed that OHCs (dotted square in Fig. 3K’’), but not IHCs, expressed Ikzf2 in *Ikzf2^V5/+^* mice at P42 (Fig. 3K-K’’). In contrast, in *Slc17a8^iCreER/^*^+^; *Tbx2^flox/-^; Ikzf2^V5/+^* (Tbx2cko-Ikzf2^V5/+^) mice at P42, besides in OHCs (dotted square in Fig. 3L’’), Ikzf2 was expressed in an additional row of cells including iOHCs (yellow arrows in Fig. 3L-L’’) and the IHCs that did not change cell fate and accordingly maintained vGlut3 expression (white arrows in Fig. 3L-L’’). It was in agreement with significant *Ikzf2* mRNA upregulation in the iOHCs (Fig. 3C and fig. S4A). Altogether, both *Ikzf2* mRNA and protein were expressed in the iOHCs. It is known that ectopic Ikzf2 is sufficient to promote IHCs to transdifferentiate into OHCs ^16,20^, thus presence of Ikzf2 protein further strengthened the ‘OHC’ features of the iOHCs.

### Tbx2 is needed in maintaining cell fate of adult cochlear IHCs

We next wondered whether Tbx2 is also required in maintaining fate of adult IHCs. The WT (n=3) and Tbx2 cko (n=3) mice were administered with tamoxifen (TMX) at P60 and P61, and analyzed at P120. Three rows of Prestin+ OHCs and one row of vGlut3+ IHCs were well aligned in WT (Fig. 4A-A’’), however, additional but discontinuous Prestin^High^/vGlut3^Low^ cells existed in the IHC region of Tbx2 cko cochleae (yellow arrows in Fig. 4B-B’’). According to our criterion above, Prestin^High^/vGlut3^Low^ cells were defined as the iOHCs and the nearby vGlut3^High^/Prestin^Low^ IHCs were suspected to be the endogenous IHCs where Tbx2 was not successfully deleted (asterisks in Fig. 4B-B’’). Quantification assay showed that 60.0 % ± 2.7 % of Prestin^High^/vGlut3^Low^ iOHCs emerged at P120 when we grouped the iOHCs in all turns together (Fig. 4C). Notably, middle turn had the least, and basal turn the most iOHCs if we counted them in different turns (Fig. 4D).

**Fig. 4.**
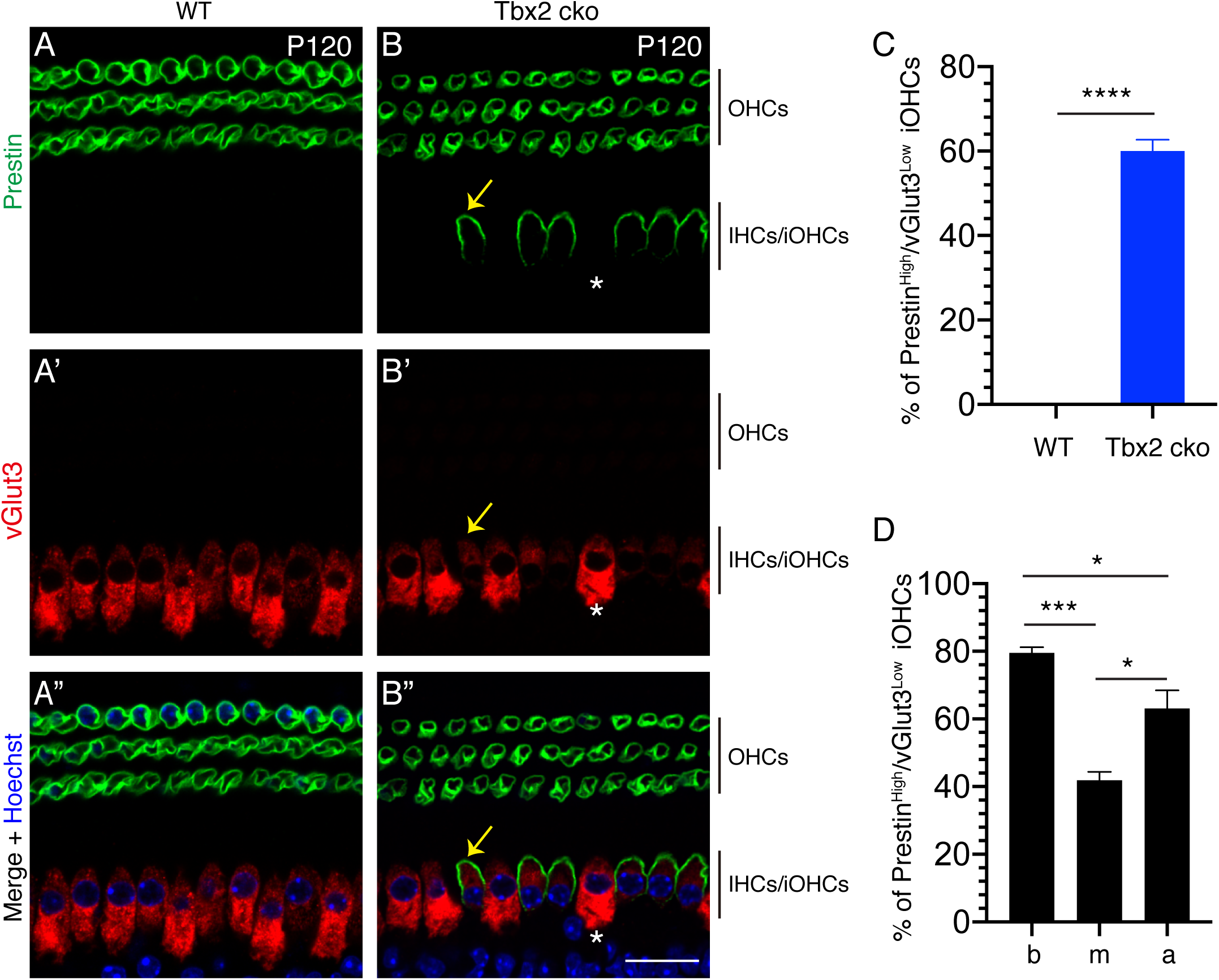
Adult IHCs also transdifferentiate into OHCs when Tbx2 is conditionally deleted. **(A-B’’)** Double staining of Prestin and vGlut3 in WT (A-A’’) and Tbx2 cko (B-B’’) cochleae. Yellow arrows in (B-B’’) mark the iOHC that expresses high Prestin but low vGlut3, whereas white asterisks label the suspected endogenous IHC that maintains high vGlut3 expression but does not express Prestin. **(C)** The averaged percentage of the Prestin^High^/vGlut3^Low^ iOHCs in all turns of Tbx2 cko cochleae. Data are presented as Mean ± SEM. **** p<0.0001. There is no iOHC in WT cochleae. **(D)** The individual percentage of the Prestin^High^/vGlut3^Low^ iOHCs in basal (b), middle (m) and apical (a) turns of Tbx2 cko cochleae. Data are presented as Mean ± SEM. * p<0.05; *** p<0.001. Scale bar: 20 μm.

Notably, vGlut3 was undetectable in the iOHCs at P42 when *Tbx2* was deleted at P2 and P3, but it was detectable, albeit faint, in the iOHCs at P120 (2 months after its deletion at P60/P61). Collectively, our data supported that Tbx2 is required in maintaining or stabilizing IHC fate at both neonatal (P2/P3) and adult ages (P60/P61). Without Tbx2, IHCs tend to transdifferentiate into OHCs and become iOHCs, and the cell fate conversion speed is faster in neonatal than the counterparts in adult IHCs.

### A new genetic model to induce temporal Atoh1 but permanent Tbx2 expression

The importance of Tbx2 in IHC development promoted us to further hypothesize that Tbx2 together with Atoh1 should be able to convert cochlear IBCs/IPhs, which are physically close to IHCs, into more differentiated new IHCs (i.e. vGlut3+) than the case via Atoh1 alone in our previous study ^18^. The IBCs/IPhs were effectively targeted by Plp1-CreER+ (Fig.5A-A’’’), as reported in other studies ^18,27,28^. In addition, we aimed to only induce temporal Atoh1 expression, mimicking the case of Atoh1 during the endogenous IHC development ^5,29–31^. Escherichia coli dihydrofolate reductase (DHFR) is the destructing domains (DDs), and fusing DHFR to both N and C termini of a protein of interest yields better temporal control effects than the case to either N or C end alone ^32^.

**Fig. 5.**
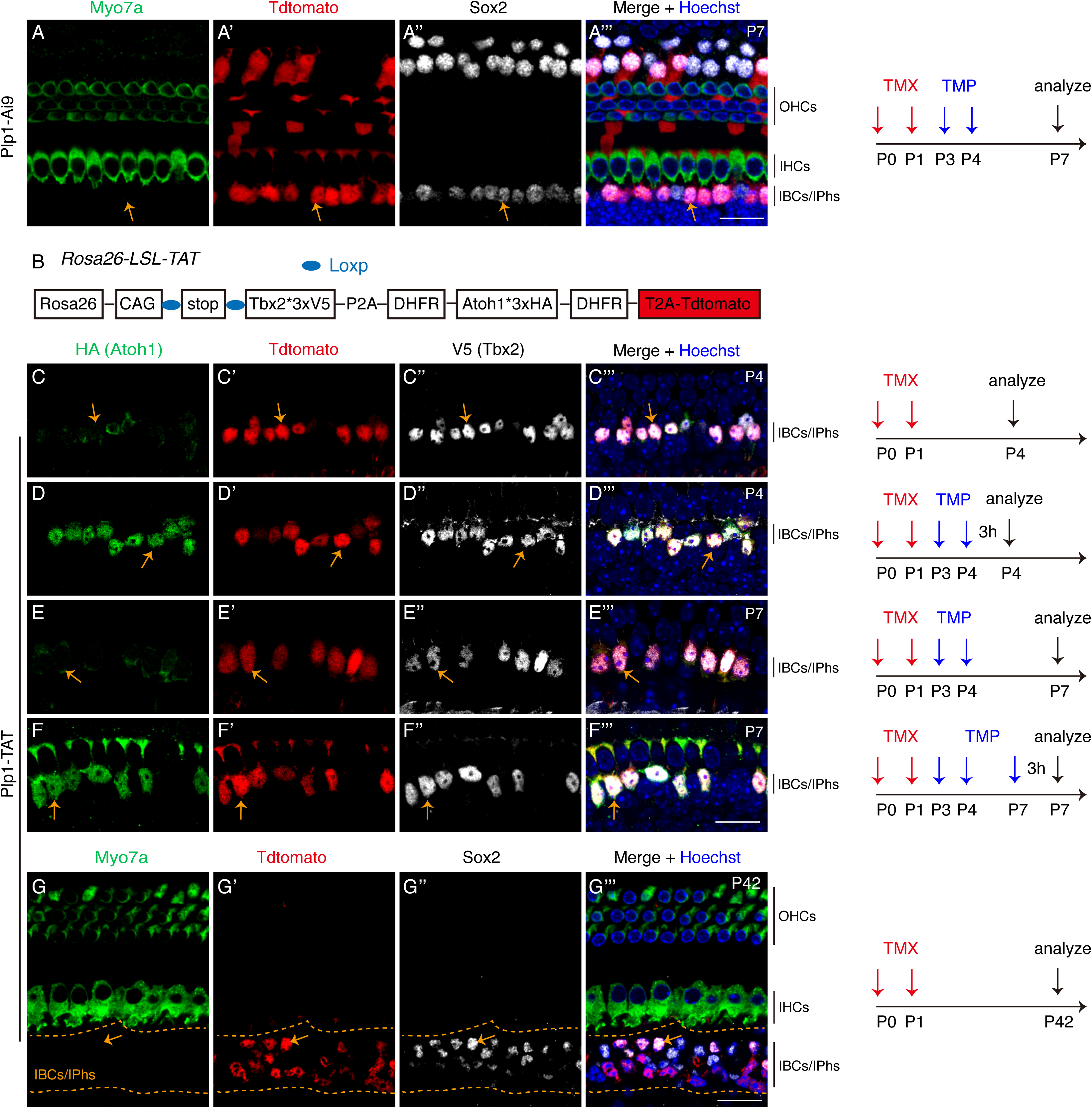
Transient Atoh1 and permanent Tbx2 ectopic expression are successfully induced in our Plp1-TAT mice. **(A-A’’’)** Triple staining of Myo7a, Tdtomato and Sox2 in control cochleae of Plp1-Ai9 mice that are administered with TMX (red arrows) and TMP (blue arrows), and analyzed at P7. Orange arrows mark one IBC/IPh that is Tdtomato+/Sox2+/Myo7a-. **(B)** The simple cartoon illustrating the genetic model to conditionally induce transient Atoh1 and permanent Tbx2. Please also refer to fig. S5 for details. **(C-F’’’)** Triple staining of HA (Atoh1), Tdtomato and V5 (Tbx2) in Plp1-TAT mice that are subjected to different treatment procedures and analyzed at different ages, as illustrated on the right sides. Orange arrows in (C-C’’’) label one IBC/IPh that expresses high protein levels of Tdtomato and Tbx2, but weak Atoh1. Differently, orange arrows in (D-D’’’) point to one IBC/IPh expressing high protein levels of Tdtomato, Tbx2 as well as Atoh1 because Atoh1 is stabilized with additional TMP treatment. Atoh1 is reversed to a weak level in Tdtomato+/Tbx2+ cells (orange arrows in E-E’’’) if mice are analyzed at P7 (3 days after TMP), but can be recovered to a high level in Tdtomato+/Tbx2+ cells (orange arrows in F-F’’’) if the third TMP is given 3 hours before final analyzing at P7. **(G-G’’’)** Triple staining of Myo7a, Tdtomato and Sox2 in cochleae of Plp1-TAT mice at P42. Arrows point to the Tdtomato+/Sox2+ cell that does not express Myo7a and is defined as IBC/IPh failing to become HCs. Scale bar: 20 μm (A’’’, F’’’ and G’’’).

Next, we constructed the *<u>Ros a26</u>*-CAG-<u>L</u>oxp-<u>S</u>top-<u>L</u>oxp-<u>T</u>bx2*3xV5-P2A-DHFR-<u>A</u>toh1*3xHA-DHFR-T2A-<u>T</u>dtomato*/+* (*Rosa26*-LSL-TAT/+) (Fig. 5B and fig. S6A-F). Thus, Atoh1 protein, which was tagged with DHFR, would be unstable and undergo rapid degradation (fig. S6G), but become temporally stable (fig. S6H), when a cell-permeable small molecule trimethoprim (TMP) is available ^33,34^. Moreover, upon Cre-mediated recombination occurs, Atoh1 (also tagged with three HA fragments), Tbx2 (tagged with three V5 fragments) and Tdtomato were transcribed from the same polycistronic mRNA.

We confirmed the transient Atoh1 and persistent Tbx2 expression in our model by four following assays. Hereafter, administration of TMX was performed at P0 and P1 and TMP was at P3 and P4, unless other ages were specifically described. First, in Plp1-CreER+; *Rosa26*-LSL-TAT/+ (abbreviated as Plp1-TAT) without TMX administration, Tdtomato+ or V5 (Tbx2) + or HA (Atoh1) + cells were never observed and data were not presented here. Secondly, Plp1-TAT mice with TMX administration were divided into two groups: No-TMP and TMP-treated. Both groups were analyzed at P4 (3 hs after the last TMP). In cochleae of no-TMP mice, we captured Tdtomato+ cells in which V5 (Tbx2) expression level was high, but HA (Atoh1) was faint or undetectable (arrows in Fig. 5C-C’’’), whereas Tdtomato+ cells expressing high Tbx2 and Atoh1 were present in TMP-treated mice (arrows in Fig. 5D-D’’’). The faint Atoh1 might be due to the occasionally incomplete degradation of Atoh1 proteins. Thirdly, while high Tbx2 and Tdtomato expression were maintained, Atoh1 expression was reversed to a faint or undetectable level, in the same cells when TMP-treated group mice were analyzed at P7 (arrows in Fig. 5E-E’’’). Fourthly, the Atoh1 level was high again if the additional (the third) TMP was administered 3 hours before sacrifice at P7 (arrows in Fig. 5F-F’’’). Collectively, we concluded that expression of Tbx2 and Tdtomato was solely TMX dependent, but Atoh1 expression was dependent on both TMX and TMP. Moreover, Atoh1 protein level was reversible and TMP treatment could only stabilize Atoh1 transiently (less than 3 days).

### Dual expression of transient Atoh1 and permanent Tbx2 successfully convert neonatal IBCs/IPhs into vGlut3+ new IHCs

We first confirmed that Tbx2 alone cannot convert neonatal IBCs/IPhs into IHCs or the general HCs by characterizing the Plp1-TAT mice that were only administered TMX and analyzed at P42. All Tdtomato+ cells, which were derived from IBCs/IPhs, failed to express Myo7a (Fig. 5G-G’’’). We did not observe Tdtomato+/vGlut3+ cells, either. Secondly, we continued to determine whether concurrent inducing high level of Atoh1 and Tbx2 would convert neonatal IBCs/IPhs into vGlut3+ new IHCs (Fig. 6A). In cochleae of control Plp1-Ai9 mice at P42 (n=3), neither IHC marker vGlut3 nor OHC marker Prestin was expressed in the Tdtomato+ cells that were IBCs/IPhs (arrows in Fig. 6B-B’’’). In contrast, Tdtomato+ cells expressing vGlut3, but not Prestin, were observed in cochleae of Plp1-TAT mice at P42 (arrows in Fig. 6C-C’’’). According to our criteria, those Tdtomato+/vGlut3+ cells were defined as new IHCs (or conservatively named as IHC-like cells). Those new IHCs were primarily adjacent to the endogenous IHCs that were vGlut3+/Tdtomato-.

**Fig. 6.**
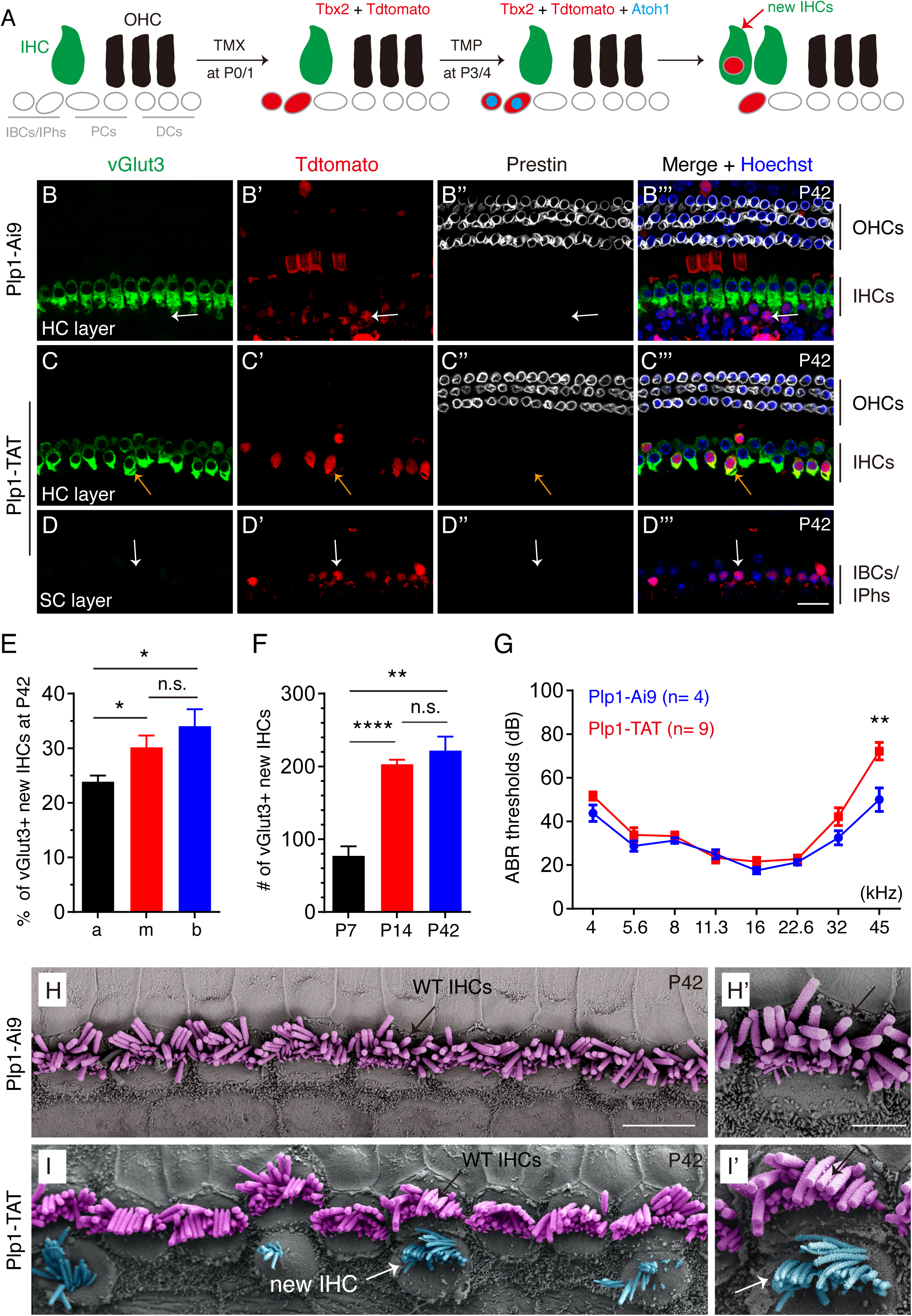
Transient Atoh1 and permanent Tbx2 convert neonatal IBCs/IPhs into vGlut3+ new IHCs. **(A)** The simple cartoon illustrating the key cellular events occurring in the reprogramming process. **(B-D’’’)** Triple staining of IHC marker vGlut3, Tdtomato, and OHC marker Prestin in control Plp1-Ai9 (B-B’’’) and experimental Plp1-TAT mice (C-D’’’) at P42. Images are visualized at HC layer (B-C’’’) or SC layer (D-D’’’). Arrows in (B-B’’’) mark one Tdtomato+ IBC/IPh cell that expresses neither vGlut3 nor Prestin. Arrows in (C-C’’’) mark one new IHC that was Tdtomato+/vGlut3+/Prestin-. Arrows in (D-D’’’) point to one Tdtomato+ IBC/IPh cell that fails to undergo cell fate change and expresses neither vGlut3 nor Prestin. **(E)** Percentage of vGlut3+ new IHCs in three cochlear turns, basal (b), middle (m) and apical turns (a), of Plp1-TAT mice at P42. Data are presented as Mean ± SEM (n=5). Percentage of new IHCs in apical is lower than middle and basal turns (* p<0.05). **(F)** Total numbers of vGlut3+ new IHCs in the entire cochleae of Plp1-TAT mice at P7 (n=3), P14 (n=5) and P42 (n=5). Data are presented as Mean ± SEM. ** p<0.01, **** p<0.0001. **(G)** ABR measurements of between Plp1-Ai9 (blue line) and Plp1-TAT (red line) mice at P42. Except the 45k Hz (*p<0.05), no statistic difference is detected. **(H-I’)** SEM analysis of control Plp1-Ai9 (H-H’) and experimental group Plp1-TAT mice (I-I’). The black arrows in (H-H’) and (I-I’) label the same endogenous IHCs, respectively, white arrows in (I-I’) mark the same new IHCs. Scale bar: 2 μm (H’); 5 μm (H); 20 μm (D’’’).

Below and adjacent to these new IHCs were Tdtomato+ cells expressing neither vGlut3 nor Prestin (arrows in Fig. 6D-D’’’), which in our study were defined as IBCs/IPhs that failed to become IHCs and were primarily located in the SC layer. Here, we only quantified the Tdtomato+/vGlut3+ new IHCs close to the endogenous IHCs. Per 200 μm cochlear duct (n=5), 8.0 ± 0.8, 9.3 ± 0.7 and 5.2 ± 0.8 new IHCs were present in basal, middle, and apical turns, respectively. Furthermore, we calculated the cell fate conversion rate by normalizing number of new IHCs to the total Tdtomato+ cells that were close to the endogenous IHCs within the same region. 34.0% ± 3.1%, 30.2% ± 2.2% and 23.9% ± 1.1% of Tdtomato+ IBCs/IPhs were respectively converted into vGlut3+ new IHCs in basal, middle and apical turns (Fig. 6E). Nonetheless, there was less than 1.5-fold difference regarding the averaged cell fate conversion rate among three turns. Thus, to simply the following analysis, we hereafter grouped three turns together and found that there were 221.8 ± 19.1 new IHCs in the entire cochleae of Plp1-TAT mice at P42 (Fig. 6F), and the averaged cell fate conversion rate was 29.5% ± 1.2%. The remaining ∼70.5% were IBCs/IPhs that failed to become new IHCs. In addition, all vGlut3+ new IHCs expressed another IHC marker Otoferlin in Plp1-TAT, but not in Plp1-Ai9 mice, at P42 (fig. S7A-B’’’). Furthermore, if using Myo7a as a marker to define the general new HCs, 239.3 ± 17.2 Tdtomato+/Myo7a+ new HCs were captured in the entire cochleae of Plp1-TAT, but not in Plp1-Ai9 mice, at P42 (arrows in fig. S7C-D’’). Due to the similar numbers (fig. S7E) and percentages (fig. S7F) of vGlut3+ new IHCs and Myo7a+ new HCs, we deduced that most, if not all, of the Myo7a+ new HCs adopted IHC fate.

Last, Auditory brainstem response (ABR) assay showed that there was no statistic difference regarding the hearing thresholds between Plp1-Ai9 and Plp1-TAT mice at P42, except the 45 kHz (Fig. 6G). It suggested that, except the high frequency (45k Hz), extra IHCs did not affect the functions of the endogenous IHCs, which was in agreement with the case in *Huwe1* mutant mice where extra IHCs are also present but the hearing ability is normal ^35^. Moreover, scanning electron microscopy (SEM) analysis showed that, relative to control Plp1-Ai9 mice which contained one row of endogenous IHCs (purple color in Fig. 6H and H’), there was an additional but discontinuous row of IHCs (blue color) whose stereocilia showed a ‘bird-wing’ pattern in Plp1-TAT mice at P42 (Fig. 6I and I’). Approximately 12.1 ± 1.2 (n = 3) extra IHCs with stereocilia were captured per 200 μm, among which 68.1% ± 4.8% contained well-organized stereocilia (white arrow in Fig. 6I). The numbers were higher than those calculated by immunostaining because we preferred to scan the area with more new IHCs with SEM. Collectively, we concluded that transient Atoh1 and permanent Tbx2 reprogrammed neonatal IBCs/IPhs into new IHCs expressing early pan-HC marker Myo7a, IHC specific markers vGlut3 and Otoferlin, as well as possessing the IHC-like stereocilia. produced by Atoh1 alone ^18^.

## Discussion

The undifferentiated cochlear sensory progenitors keep proliferating until E14.5, followed by a differentiation wave in a basal to apical gradient ^36,37^. Atoh1 is absolutely essential in specifying the general HC fate, as both IHCs and OHCs are lost in *Atoh1*^-/-^ mice^4^. Recently, two TFs, Insm1 and Ikzf2, are reported to be critical for OHC development, as their mutants exhibit similar OHC phenotypes ^16,17^. Then, what is the essential gene for IHC development? We hypothesized that it should also be a TF protein that is highly expressed in IHCs, but not OHCs. Our study showed that Tbx2, which indeed is a TF protein, is uniformly expressed in all undifferentiated sensory progenitors by E13.5, but afterwards is gradually repressed in the lateral cochlear progenitors that would develop into OHCs and the nearby SCs (fig. S1). Notably, the expression of Tbx2 is highly and persistently maintained in IHCs. It is known that ablation of Tbx2 at early embryonic ages leads to cochlear hypoplasia ^23^. Therefore, we chose to conditionally and specifically delete *Tbx2* in neonatal IHCs that are immature and undergo active differentiation, by which the phenotypes are clean and easy to explain. Overall, our data demonstrated that the neonatal IHCs, upon losing Tbx2, gradually decrease vGlut3, Otoferlin or Slc7a14, and conversely increase Prestin. As expected, the final transcriptomic profiles of those *Tbx2 ^-/-^* IHCs are similar to the endogenous OHCs, which is why we defined those IHCs as iOHCs.

Is Ikzf2 protein derepressed in these iOHCs? It is a question of importance because Ikzf2 alone can make IHCs behave like OHCs ^16,20^. *Ikzf2* mRNA indeed is expressed in the iOHCs (Fig. 3). We speculated that Tbx2 promotes IHC fates partially by repressing Ikzf2 expression. However, we cannot rule out the possibility that it is a secondary effect following cell fate changes. Whether Tbx2 directly binds *Ikzf2* cis-regulatory elements warrants future *in vivo*Tbx2 CUT&RUN assays in cochlear tissues.

How to regenerate functional cochlear IHCs from non-sensory SCs is a long-term goal for inner ear biologists. Previous new IHCs regenerated from neonatal IBCs/IPhs via Atoh1 ectopic expression only express nascent HC marker Myo6 or Myo7a, and fail to turn on late IHC marker vGlut3 ^18^. Two possible interpretations are proposed: 1) Atoh1 is transiently expressed in wild type IHCs, but it is persistently induced in the immature new IHCs ^18^; 2) Other key genes are needed to further drive differentiation of the new IHCs. In our current study, we designed a new genetic approach by which transient Atoh1 expression is induced as well as persistent Tbx2 expression is effectively driven. Excitingly, we found that the differentiation status is much more advanced than previously reported ^18^. Furthermore, the reprogramming efficiency by Atoh1 and Tbx2 is 29.5%, which was higher than the 17.8% by Atoh1 alone ^18^. Collectively, we proposed that synergistic interactions between Atoh1 and Tbx2 are present during the cell fate conversion process, as the case between Atoh1 and Ikzf2 in our OHC regeneration study ^20^, or between Atoh1 and Pou4f3 during the endogenous HC development ^38^. However, the puzzle is why the neonatal IBCs/IPhs that express the endogenous Tbx2 cannot become vGlut3+ new IHCs when ectopic Atoh1 is induced ^18^. We further speculated that Tbx2 has a dose-dependent effect on cell fate determination.

Last but not the least, besides in IHCs, Tbx2 is also highly expressed non-sensory SCs that are distributed in GER region in neonatal cochleae, including the IBCs/IPhs (Fig. 1). It is known that those SCs in GER regions are plastic and can replenish themselves after damage at neonatal, but not adult cochleae ^28,39^. In agreement with normal Tbx2 in the SCs, further ectopic Tbx2 cannot convert IBCs/IPhs into new HCs or new IHCs. Because Atoh1 alone does reprogram neonatal IBCs/IPhs into immature IHCs ^18^, we speculated that Tbx2 entitles IBCs/IPhs to have intrinsic properties to become IHCs rather than OHCs. Alternatively, if cell change occurs, Tbx2 prevents IBCs/IPhs from transdifferentiating into OHCs, reminiscent of how Tbx2 represses expression of OHC genes in the endogenous IHCs. Nonetheless, the exact roles played by Tbx2 in those SCs warrants future studies. Finally, while we were finishing the studies and in the process of submission, we were aware of a similar study where the first part of our study, that is the essential role of Tbx2 in normal cochlear IHC development, was just reported ^40^. Although with different genetic approaches, similar conclusion was drawn. The two independent studies thus highlighted the critical roles of Tbx2 in IHC fate specification and maintenance.

## Materials and Methods

### Mouse models

The mouse strains Plp1-CreER+ (Stock# *005975), Rosa26*-LSL-Tdtomato/+ (Ai9, Stock#: 007909) were from The Jackson Laboratory*. Slc17a8*-P2A-iCreER/+, which is also known as *vGlut3-P2A-iCreER/+,* is reported and described in details in our previous study ^19^. Both male and female mice were used. Tamoxifen (TMX, Cat# T5648, Sigma-Aldrich) was dissolved in corn oil (Cat# C8267, Sigma-Aldrich) and was administrated at 3mg/40g body weight. Trimethoprim (TMP, Cat# T0667; Sigma-Aldrich) was dissolved in 1×phosphate buffer saline (PBS) and TMP was administered at 300 μg/g body weight. All mice were bred and raised in SPF-level animal rooms, and animal procedures were performed according to the guidelines (NA-032-2019) of the IACUC of Institute of Neuroscience (ION), CAS Center for Excellence in Brain Science and Intelligence Technology, Chinese Academy of Sciences.

### Generation of Tbx2-*HA/+*, Tbx2^+/-^, Tbx2^flox/+^, Ikzf2^V5^/+ *and* Rosa26-*LSL-TAT/+ mouse strains*

First, the Tbx2-HA+ was constructed by Crispr/Cas9-mediated homologous recombination in one cell stage of mouse zygotes illustrated in Fig. S1. The sgRNA close to the TGA stop codon is: 5’-GACCCCCGACCTGCCACGCG -3’. Secondly, Tbx2^+/-^ was generated by injecting two pre-tested efficient sgRNAs (sgRNA-1 and 2 in Fig. S2) with Cas9 mRNA. The two sgRNA sequences are below: *5’-*TGACCCGCCGTAAGGGCCTG- 3’, and 5’-AGGCTCCGAGGCGCCGACGT-3’. Thirdly, *Tbx2*^flox/+^ was also produced by the Crispr/Cas9 approach. Two sgRNAs on the left and right sides of *Tbx2* exon2, *Cas9* mRNA and the targeting vector (Fig. S2E) together were injected to one cell stage of mouse zygotes. The two sgRNA sequences (Fig. S2D) are *5’-*ATGTCATGTCATTGTCGGGA-3’ and 5’-AGGGGCCCCCACAGTCGAGT-3’. The *Ikzf2^V5^/+* was generated via the Crispr/Cas9 approach, as illustrated in Fig. S5. The sgRNA close to the TAG stop codon is 5’-AGGGGAGCACACATTCCACT-3’. Lastly, *Rosa26*-LSL-TAT/+ was similarly constructed by the Crispr/Cas9 approach, according to the design described in Fig. S6. The sgRNA used in *Rosa26* locus is *5ʹ-*ACTCCAGTCTTTCTAGAAGA-3ʹ.

For all mouse strains above, once the Founder 0 (F0) mice were born, they were screened by tail PCR. The F0 mice with potentially correct gene targeting were chosen and crossed with C57BL/6 wild type mice to produce the germ line stable F1 mice that were subjected to the second round of tail PCR screening. Except the *Tbx2^+/-^*, all other strains were subjected to Southern blot. Only the F1 mice without random insertion of the targeting vectors were chosen for further breeding. Notably, PCR primers that were used for genotyping each strain and their amplicon sizes were described in table S3.

### Sample processing and immunofluorescence assays

The mice were perfused with fresh ice-cold 1xPBS and 4% paraformaldehyde (PFA) in 1xPBS (PBS, pH 7.4) after anesthetization. Then, the inner ear samples were carefully dissected and further fixed in 4% PFA solution on rotator overnight at 4°C. On the next day, after three times 1xPBS wash, the inner ears were further decalcified in 120 mM EDTA (Cat# ST066, Beyotime) in 1x PBS at room temperature for 24 hours at the ages range from P7 to P14, or for 48 hours over P14. Inner ear samples were further washed three times with 1x PBS before proceeding for following whole mount dissection. Inner ear samples younger than P7 were dissected without EDTA treatment.

The cochlear ducts were divided into three pieces, basal, middle and apical turns. Each turn was subjected to immunostaining in parallel but in different tubes. Briefly, cochlear samples were first incubated by blocking solution containing 1% Triton X-100 (X100-500ML, Sigma) and 5% Bovine Serum Albumin (BSA, Cat#: BP1605, Fisher Scientific), followed by incubation with primary antibodies dissolved in solutions containing 0.1% Triton X-100 and 5% BSA on rotator overnight at 4°C. On the second day, samples were washed three times with 0.1% Triton X-100 dissolved in 1xPBS, and then incubated in the solution containing the secondary antibody, 0.1% Triton X-100 and 5% BSA.

The following primary antibodies were used in this study: anti-HA (rat, 1:200, 11867423001, Roche), anti-V5 (mouse, 1:500, MCA1360, Bio-rad), anti-Prestin (goat, 1:1000, sc-22692, Santa Cruz), anti-vGlut3 (rabbit, 1:500, 135203, Synaptic Systems), anti-Otoferlin (mouse, 1:500, ab53233, Abcam), anti-Myo7a (rabbit, 1:500, 25-6790, Proteus Bioscience), anti-Ctbp2 (mouse, 1:200, 612044, BD Biosciences), anti-Sox2 (goat, 1:1000, sc-17320, Santa Cruz), anti-Myo6 (rabbit, 1:500, 25-6791, proteus-biosciences), anti-Slc7a14 (rabbit, 1:500, HPA045929, Sigma-Aldrich). Finally, samples were counterstained with Hoechst 33342 solution in PBST (1:1,000, 62249, Thermo Scientific) to visualize nuclei, and were mounted using Prolong Gold antifade medium (P36930, Thermo Scientific). The detailed immunostaining protocols have been described previously ^41^. Nikon C2 (Nikon, Japan), TiE-A1 plus (Nikon, Japan), or NiE-A1 plus (Nikon, Japan) confocal microscopes was used to capture images.

### Calculating percentages of the iOHCs, ribbon synapses and the new IHCs

Three pieces of the cochlear duct were first scanned under 10x lens of confocal microscope. The total length of the three pieces was calculated by drawing a line between IHCs and OHCs via Image J software. Then, each cochlea was divided into three portions with equal length, by which each region was assigned into basal or middle or apical turns. In the Tbx2cko model, numbers of total IHCs (or cell types derived from the endogenous IHCs) were calculated by adding the iOHCs (Prestin^High^/vGlut3^Low^, Prestin^High^/Otoferlin^Low^, or Prestin^High^/Slc7a14^Low^) and the IHCs failing to become iOHCs (Prestin^High^/vGlut3^Low^, Prestin^Low^/Otoferlin^High^ or Prestin^Low^/Slc7a14^High^). We scanned the entire cochlear turns by confocal microscopes (60x lens) in order to quantify the percentages of the iOHCs (Figures 2A-H, Fig. S3A-F and Fig. 4A-D) with minimal variation. The percentage of iOHCs was calculated by normalizing numbers of all iOHCs to the total IHCs. In terms of quantifying the numbers of Ctbp2+ puncta that represent the ribbon synapse, we chose the 16 kHz frequency region based on the method mentioned previously ^42^ and scanned the cochlear samples with 0.41 µm interval during the z-stack scanning (Fig. S3G-I).

For quantifying numbers or percentages of the vGlut3+/Tdtomato+ new IHCs (Fig. 6C-F) or Myo7a+/Tdtomato+ nascent HCs (Fig. S7), we scanned three areas (60x) in each turn, by which an average number was obtained per mouse. Percentages of vGlut3+ new IHCs or Myo7a+ nascent HCs were calculated by normalizing numbers of the vGlut3+/Tdtomato+ or Myo7a+/Tdtomato+ to the total Tdtomato+ cells (but close to IHCs only) that included new IHCs/HCs and the Tdtomato+ IBCs/IPhs failing to become new IHCs or HCs. Notably, the total numbers of vGlut3+ new IHCs (Fig. 6F) or Myo7a+ (Fig. S7E) in the entire cochlear duct were calculated by multiplying the numbers under confocal microscope (60x) with ∼5733.3 μm (total cochlear length) / ∼230.9 μm (length in 60x lens). Statistic analysis was performed via one-way ANOVA and student’t test with Bonfeerroni corrections were used.

### ABR measurement and SEM preparation and analysis

ABR were measured at 4k, 5.6k, 8k, 11.3k, 16k, 22.6k, 32k and 45k Hz at P42, following our previously published protocol ^19^. Student’s t-tests were applied to determine the statistical significance regarding the hearing thresholds of the same frequency among different mice (Figures 2I and 6G). We followed the same SEM protocol that was described in details in our previous study ^20^.

### Preparation of cell suspensions and smart-seq single-cell RNA-Seq

The cochlear samples from Slc17a8-Ai9 at P14 or P30, and Slc17a8-Tbx2cko-Ai9 at P30 were dissected out and neural regions were removed as much as possible, followed by incubation in choline chloride solution containing 20 U/mL papain (Cat# LK003178, Worthington) and 100 U/mL DNase I (Cat# LK003172, Worthington) for 20 min at 37°C, and this was followed by digestion with a protease (Cat# P5147, Sigma; 1 mg/mL) and dispase (Cat# LS02104, Worthington; 1 mg/mL) for 20 min at 25°C. The components of the choline chloride solution contained 92 mM choline chloride, 2.5 mM KCl, 1.2 mM NaH_2_PO_4_, 30 mM NaHCO_3_, 20 mM HEPES, 25 mM glucose, 5 mM sodium ascorbate, 2 mM thiourea, 3 mM sodium pyruvate, 10 mM MgSO_4_.7H_2_O, 0.5 mM CaCl_2_.2H_2_O, and 12 mM N-acetyl-L-cysteine. Next, the post-digested cochlear samples were gently triturated using fire-polished glass Pasteur pipettes (13-678-20b; Fisher) with 4 different pore sizes (largest size the first, and smallest one the last).

Finally, the dissociated Tdtomato+ cells were manually picked under the fluorescence dissecting microscope (M205FA, Leica). Notably, in the Slc17a8-Tbx2cko-Ai9 model we could distinguish the Tdtomato+ iOHCs (smaller size, picked) with the tdtomato+ IHCs that did not undergo cell fate change (larger size, not picked). The picked endogenous IHCs and iOHCs were immediately subjected to reverse-transcription and cDNA amplification using a Smart-Seq HT kit (Cat# 634437, Takara). The post-amplified cDNA (1 ng) of each cell was processed for final library construction by TruePrep DNA Library Prep Kit V2 for Illumina (Cat# TD503-02, Vazyme) and a TruePrep Index Kit V2 for Illumina (Cat# TD202, Vazyme). The final libraries were subjected to paired-end sequencing on the Illumina Novaseq platform, yielding ∼4G raw data per library.

### Bioinformatics analysis

The FASTQ files of smart-seq data were aligned to the mouse genome (GRCm38 mm10) using Hisat2 alignment package (v2.1.0) ^43^. Raw count matrices were generated by HTseq (v0.10.0) ^44^, and the transcript per million (TPM) values were calculated via StringTie (v1.3.5) ^45^. For integration analysis of 10x Genomics and Smart-seq data, Normalization (NormalizeData), principal component analysis (RunPCA), dimensional reduction (RunUMAP), unsupervised clustering (FindNeighbors/FindClusters) and integration (FindIntegrationAnchors/IntegrateData) were carried out by Seurat (v3.2.3) ^46^. The differentially (p < 0.05, absolute value of (log2 Fold Change) > 2) expresses genes (DEGs) between P30_WT IHCs and P30_WT OHCs, as well as between P14_WT IHCs and P14_iOHCs, were calculated via DESeq2 (v1.34.0) ^47^. Furthermore, DEGs between P30_WT OHCs and P30_WT IHCs with the averaged TPM >16 were defined as OHC and IHC genes.

For 10x genomic single cell data from the previous study ^26^, cells expressing above zero of *Insm1, Myo6,* and *Atoh1* were identified as E16_WT OHCs. P1_WT OHCs were defined as cells with above zero expression of *Bcl11b, Myo6, Myo7a,* and *Atoh1;* P7_WT OHCs were classified as those in which the expression level of *Slc26a5, Myo6, Ocm,* and *Ikzf2* were all above zero. Similarly, E16_WT IHCs and P7_WT IHCs were defined as cells in which the expression levels of *Myo6,* and *Fgf8* were all above zero. Cells expressing above zero level of *Myo7a, Myo6* and *Fgf8* were classified as P1_WT IHCs.

Trajectory analysis was performed in Monocle (v2.14.0) ^48^. The pre-processed matrix in the Seurat object above were used. The top 2000 most variable genes assessed from the Seurat object (“FindVariableFeatures”) were used as input to simulate the pseudotime trajectory. Cells were fitted onto the backbone of the trajectory graph by using the “orderCells” function in Monocle. All the raw data of our single-cell RNA-seq analyses have been deposited in the GEO (Gene Expression Omnibus) under the accession number GEO: GSE199369.

## Author contributions

Z. Bi and X. Li contributed to constructions of the new mouse models, mouse breeding, sample preparation of immunostaining and SEM, data analysis, single cell manual picking and preparing figures, method part of the manuscript; M. Ren contributed to bioinformatic analysis; Y. Gu contributed to southern blot assay; T. Zhu contributed to SEM analysis; S. Li, S. Sun and Y. Sun also contributed to single cell manual picking; S. Li generated the *Ikzf2^V5/+^* mice. G. Wang contributed to mouse zygote injection; Z. Liu contributed to the overall design of the project, supervision, manuscript writing and funding support.

## Materials & Correspondence

Material requests should be addressed to Dr. Zhiyong Liu, the correspondence author of this manuscript via.

## Supplemental Figure Legends

**Fig. S1.**
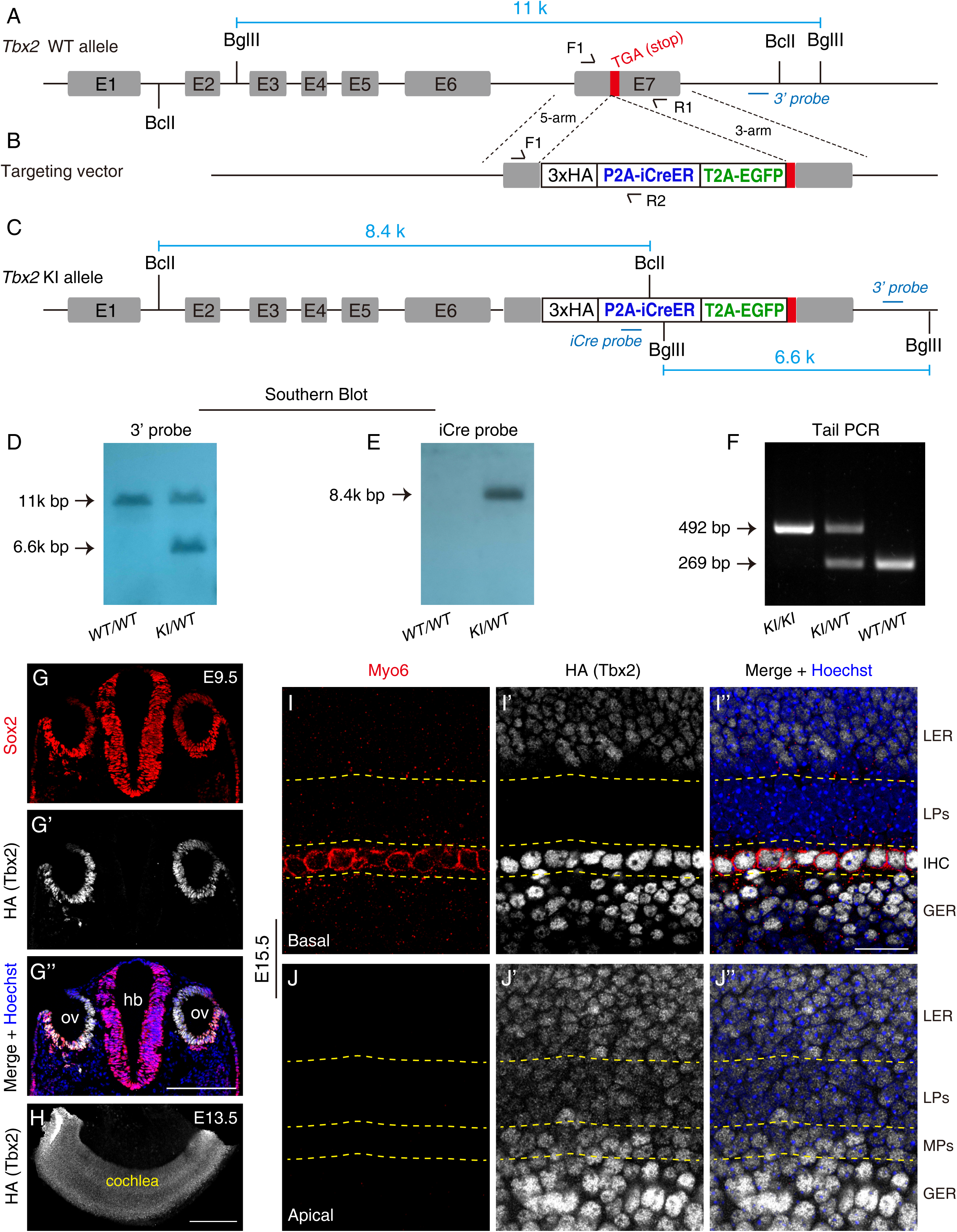
Tbx2 is turned on in early otocyst cells but its expression gradually is depressed in cochlear lateral progenitors. **(A-C)** The detailed illustration of how Tbx2*3xHA-P2A-iCreER-T2A-EGFP/+ (*Tbx2*-HA/+) is designed. Immediately before the stop codon in Tbx2 WT allele (A), a segment containing 3xHA-P2A-iCreER-T2A-EGFP (B) is inserted and the final targeted Tbx2 allele is shown in (C). **(D-E)** Southern blot results via both 3’-probe (D) and internal iCre probe (E) confirm the absence of random insertion of the donor DNA (B). **(F)** The representative tail PCR image that is able to distinguish knockin (KI) and WT alleles. **(G-G’’)** Dual staining of HA (Tbx2) and Sox2 in *Tbx2*-HA/+ embryos at E9.5. Tbx2 is highly expressed in otic vesicle (OV) cells. **(H)** The whole mount cochlear sample of *Tbx2*-HA/+ is stained by HA antibody. Tbx2 is broadly expressed in cochlear cells. **(I-J’’)** Double staining of Myo6 and HA (Tbx2) in basal (I-I’’) and apical (J-J’’) turns of *Tbx2*-HA/+ cochlear samples at E15.5. Tbx2 is maintained in apical, but becomes undetectable in basal, lateral progenitors (LPs). Scale bar: 200 μm (G’’ and H); 20 μm (I’’).

**Fig. S2.**
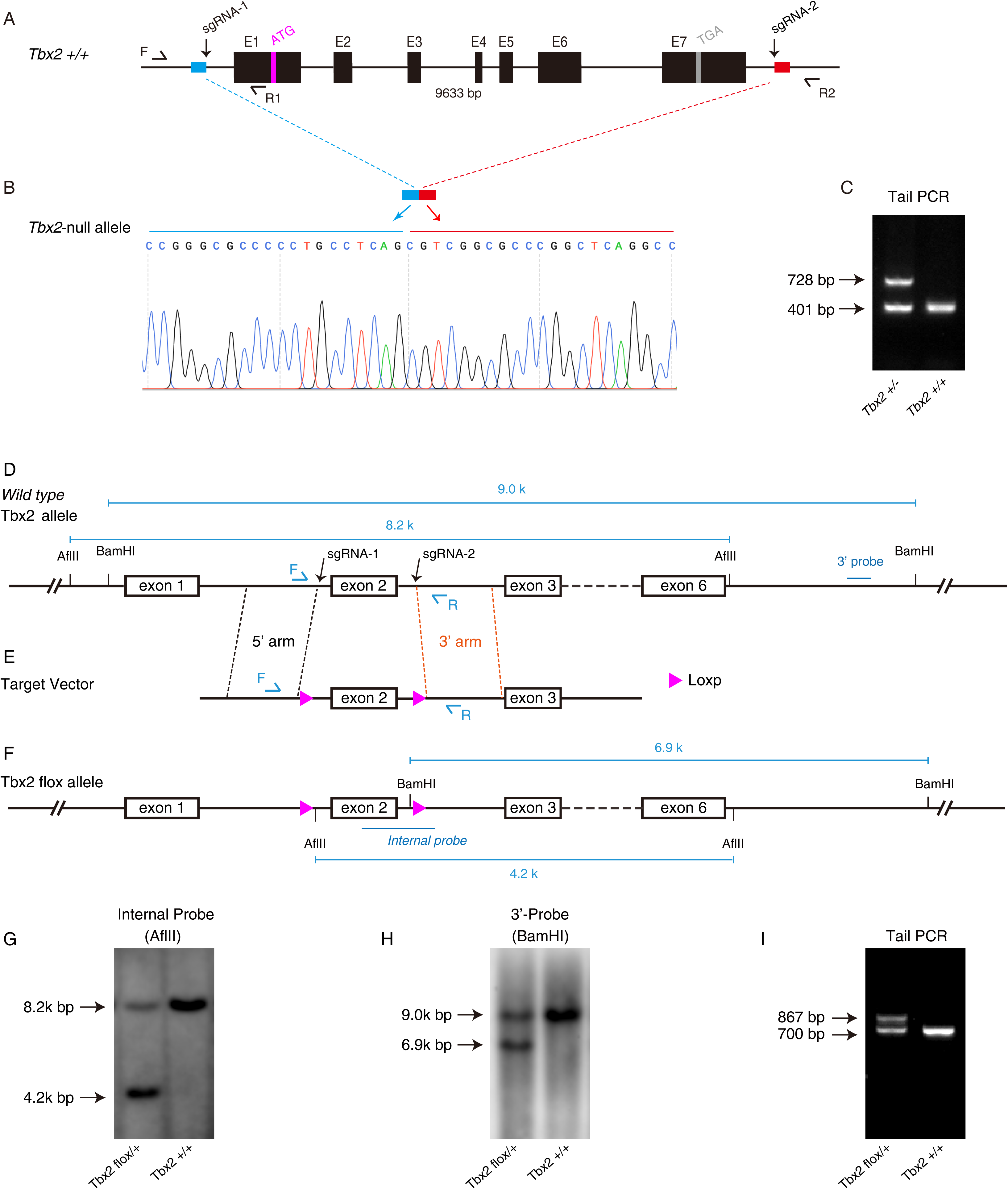
Generation of germ line *Tbx2^+/-^* and conditional *Tbx2 ^flox/+^* strains. **(A-C)** Tbx2 exons and introns between the sgRNA-1 and sgRNA-2 (arrows in A) are deleted and confirmed by Sanger sequencing (B). Wild type and Tbx2-null alleles show tail PCR bands with 401 bp and 728 bp, respectively (C). **(D-F)** The *Tbx2* WT allele (D) is recombined with the targeting vector in which the exon 2 is flanked by containing two loxp sequences (B), producing the post-targeted floxed *Tbx2* allele (F). **(G-H)** Southern blot results via both the internal probe (G) and 3’-probe (H) show the absence of random insertion of the targeting vector. **(I)** The representative tail PCR image that is competent to identify WT and floxed *Tbx2* alleles.

**Fig. S3.**
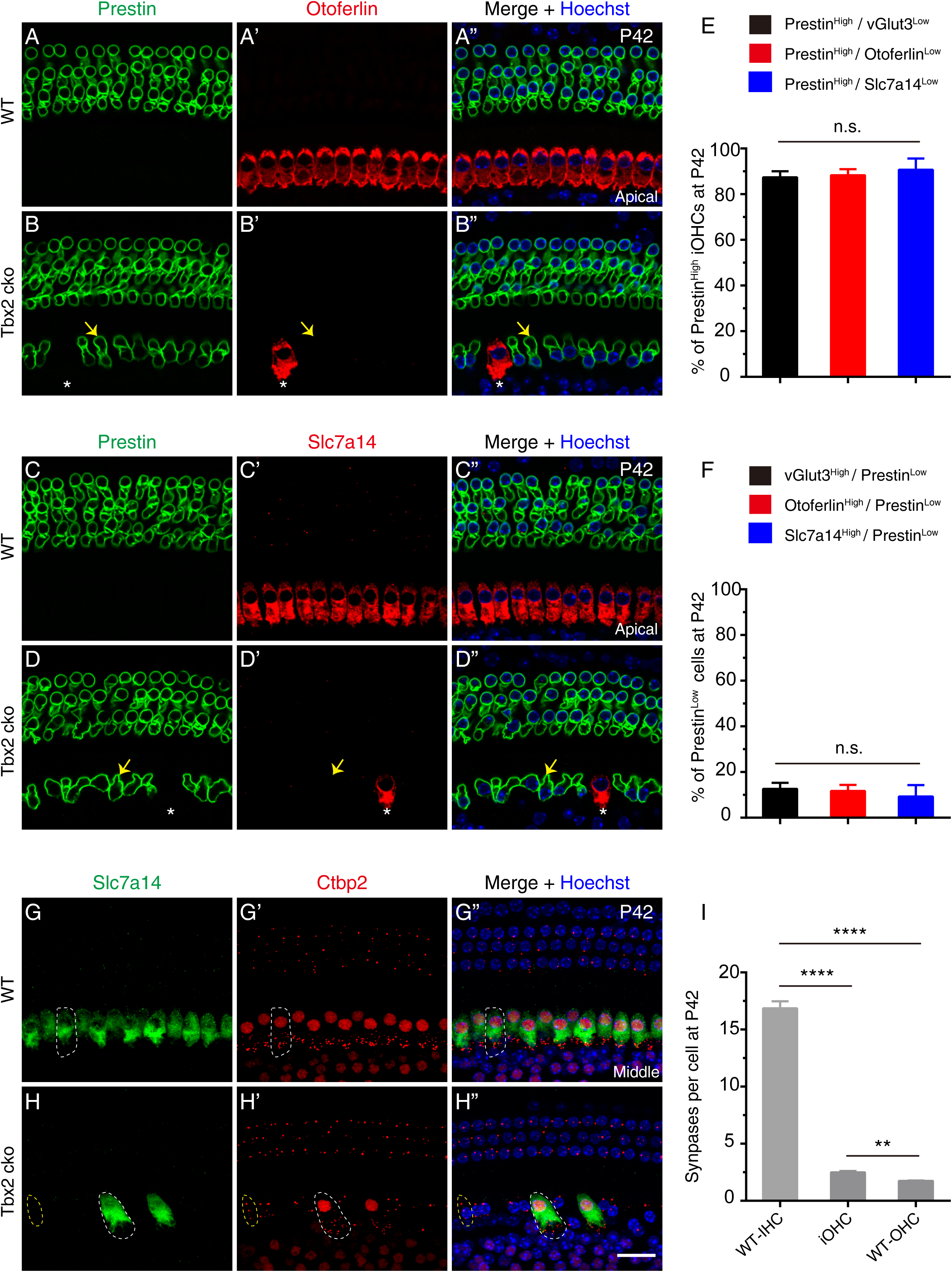
IHC markers Otoferlin and Slc7a14 as well as number of ribbon synapse are decreased in the absent of Tbx2 at P42. **(A-D’’)** Double staining of IHC marker Otoferlin (A-B’’) or Slc7a14 (C-D’’) and Prestin in WT (A-A’’ and C-C’’) and Tbx2 cko (B-B’’ and D-D’’) cochlear samples. Yellow arrows label one Prestin^High^/Otoferlin^Low^ (B-B’’) and one Prestin^High^/Slc7a14^Low^ (D-D’’) iOHC, respectively, whereas the white asterisks represent the IHCs fails to undergo cell fate conversion and maintain Otoferlin (B-B’’) or Slc7a14 (D-D’’), which are respectively included into the Otoferlin^High^/Prestin^Low^ or Slc7a14^High^/Prestin^Low^ population. **(E-F)** Quantification of the Prestin^High^/vGlut3^Low^, Prestin^High^/Otoferlin^Low^ and Prestin^High^/Slc7a14^Low^ (E) and the vGlut3^High^/Prestin^Low^, Otoferlin^High^/Prestin^Low^ or Slc7a14^High^/Prestin^Low^ (F). No significant difference is detected. **(G-H’’)** Double staining of Slc7a14 and ribbon synapse marker Ctbp2 in WT (G-G’’) and Tbx2 cko (H-H’’) cochlear samples. White dotted circles label one endogenous IHC in WT cochleae (G-G’’) or one IHC that do not undergo cell fate change and maintain Slc7a14 expression (H-H’’). Yellow dotted circles mark one iOHC. **(I)** Quantification of synapse numbers per IHC or iOHCs or WT-OHC, via numbers of Ctbp2+ puncta. Data are presented as Mean ± SEM. ** p<0.01, **** p<0.0001. The synapse numbers in iOHCs are less than WT-IHC, but more than WT-OHCs. Scale bar: 20 μm.

**Fig. S4.**
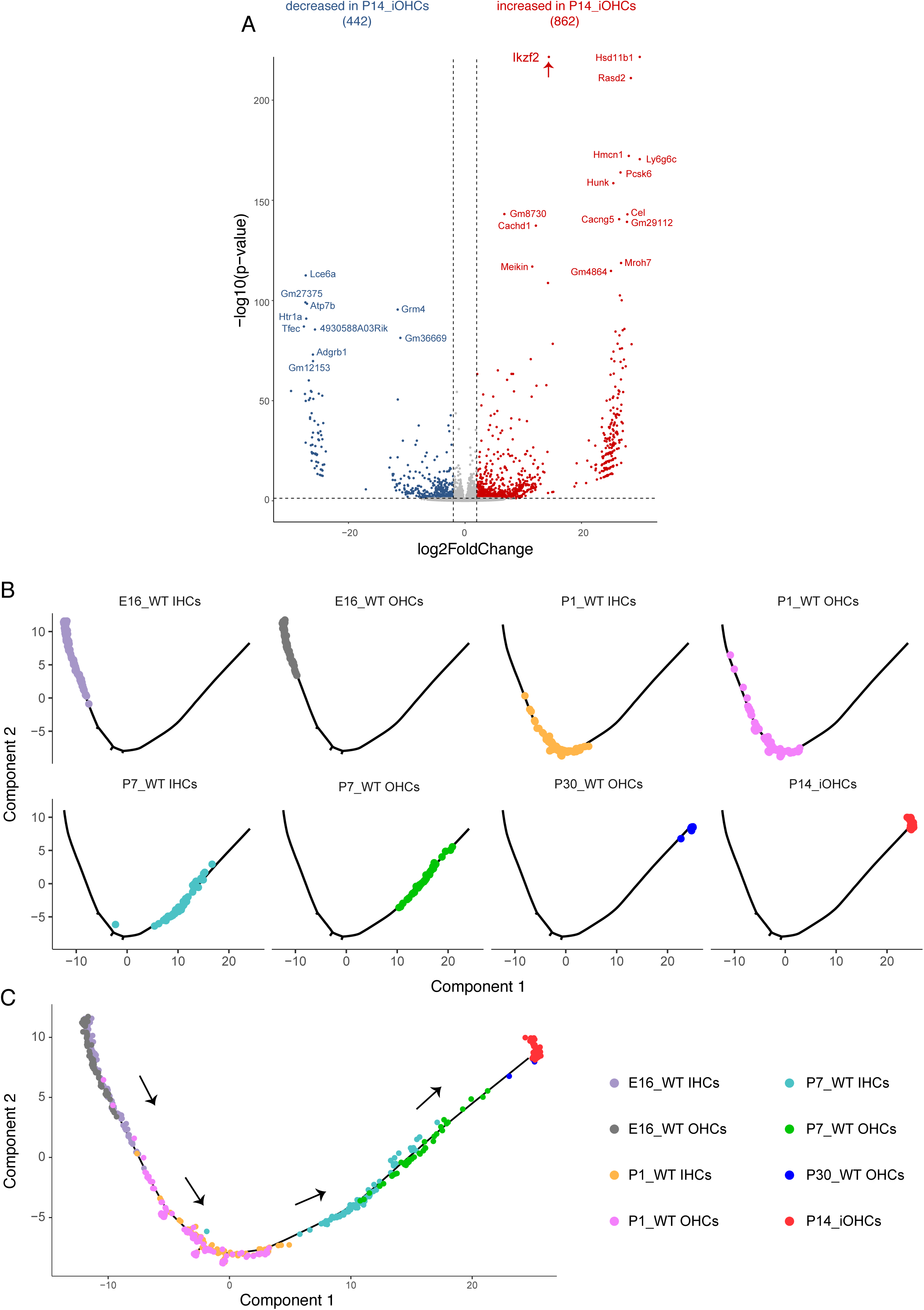
Molecular features of the P14_iOHCs. **(A)** Volcano plot of the all differentially expressed genes between P14_WT IHCs and P14_iOHCs. Volcano plot is according to gene fold change and p-value of statistics between P14_WT IHCs and P14_iOHCs. However, the absolute expression level of each gene is not considered. Red arrow marks the *Ikzf2* gene that is also highlighted in Fig. 4C where gene expression level is considered. **(B-C)** Trajectory analysis of mixed cell population that include 8 cell types: E16_WT IHCs, E16_WT OHCs, P1_WT IHCs, P1_WT OHCs, P7_WT IHCs, P7_WT OHCs, P30_WT OHCs, and the P14_iOHCs. They are plotted separately in (B) as well as presented together in (C). The size of (C) is larger than each panel in (B) in order to better visualize the distribution of each cell type. Arrows in (C) represent the calculated developmental direction.

**Fig. S5.**
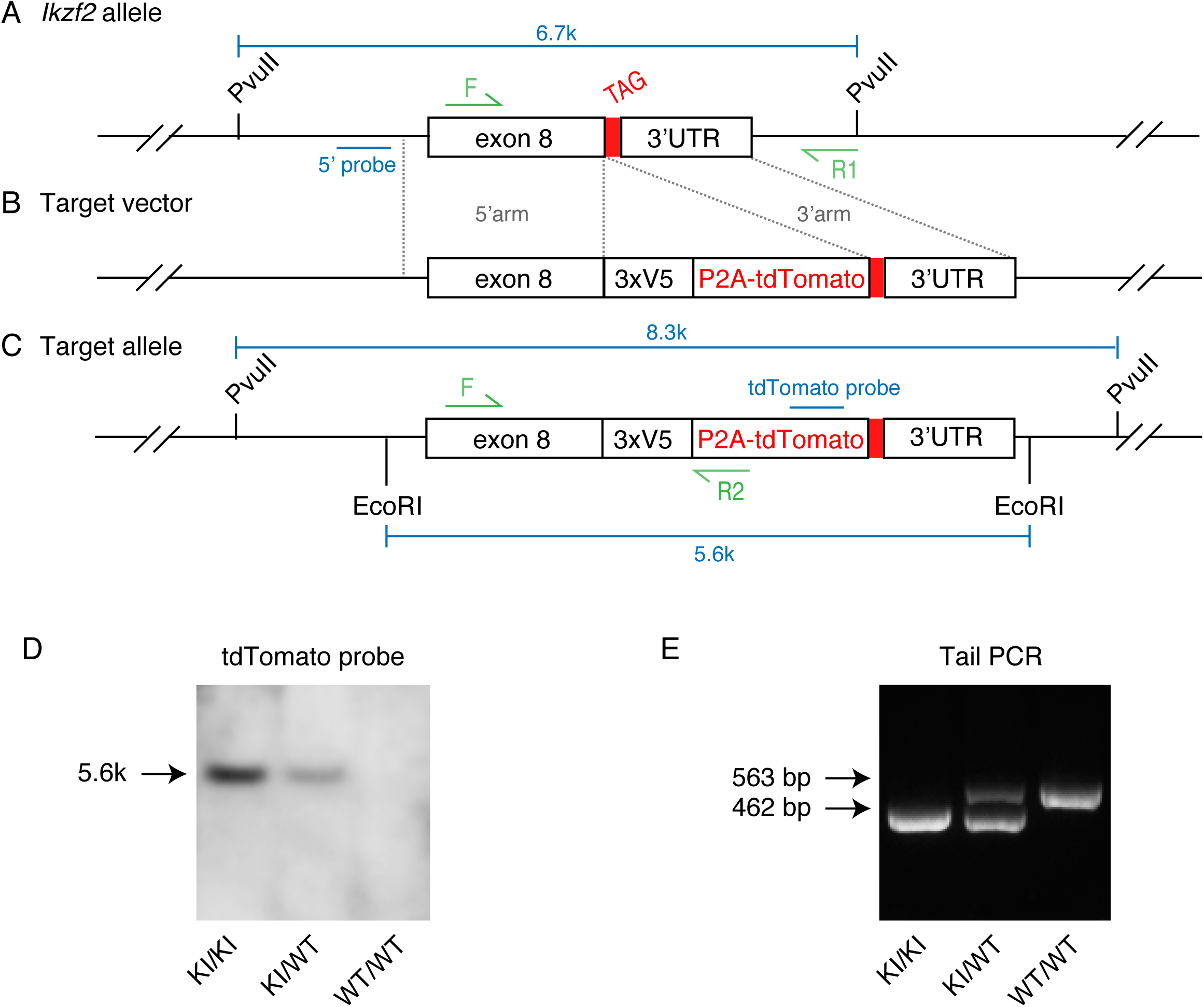
Generation of *Ikzf2*\*3xV5-P2A-Tdtomato/+ (Ikzf2^V5/+^) mouse strain. **(A-C)** Immediately prior to the TAG stop codon in WT *Ikzf2* allele (A), a fragment comprising of 3xV5-P2A-Tdtomato (B) is inserted, generating a post-targeted allele (C). **(D)** Southern blot with internal Tdtomato probe yields a single 5.6k bp band, confirming the absence of random insertion of the target vector (B). **(E)** The representative tail PCR gel image that is used to distinguish WT and the knockin (KI) alleles.

**Fig. S6.**
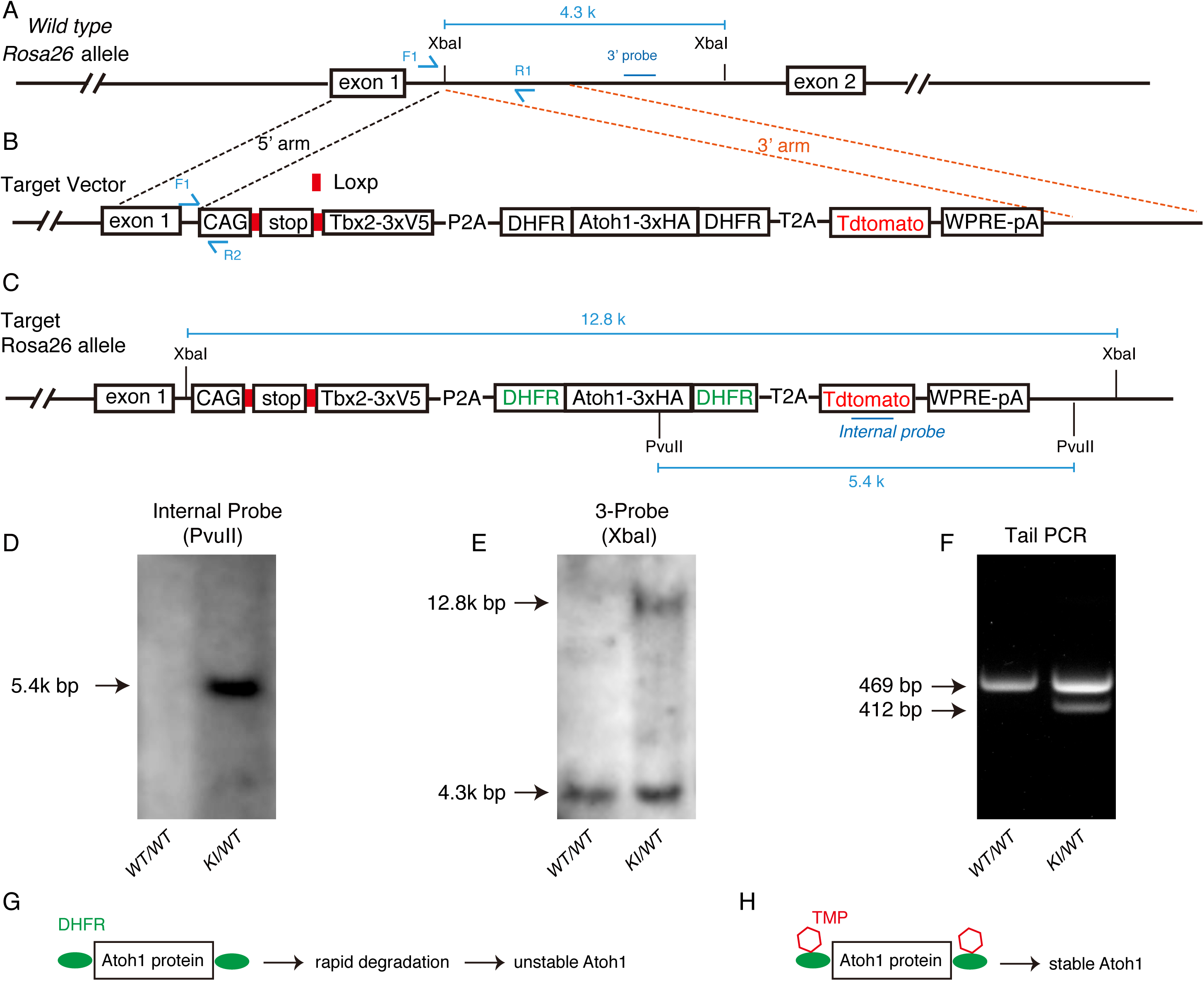
Construction of the new *Rosa26-LSL-TAT/+*. **(A-C)** In wild type *Rosa26* allele (A), we insert a long polycistronic DNA element containing Tbx2 that is fused with three V5 fragments at its C-terminus, and Atoh1 that is fused with one DHFR on its N-terminus, and three HA fragment and the other DHFR element, and Tdtomato. The final post-gene target allele is listed in (C). **(D-E)** Southern blot of internal (D) and external 3’ (E) probe are performed to confirm absence of random insertion of targeting vector in *Rosa26-LSL-TAT*/+ mouse genome. **(F)** Tail DNA is readily to distinguish wild type (469 bp) and knockin (KI, 412 bp) alleles. **(G and H)** Schematic diagram of stability of Atoh1-DHFR cassette relying on the existence of TMP.

**Fig. S7.**
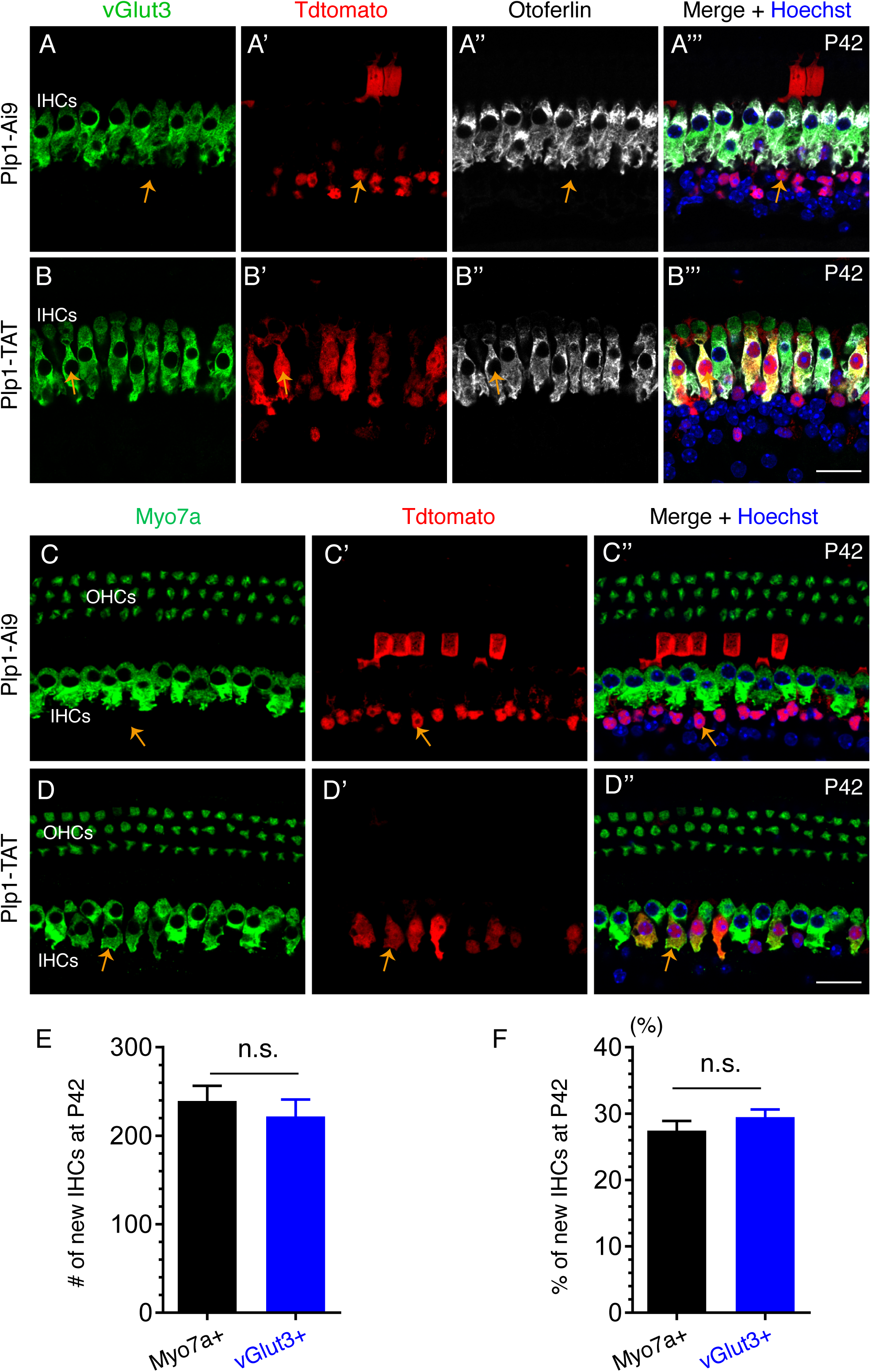
New IHCs also express another IHC marker Otoferlin and pan-HC marker Myo7a. **(A-B’’’)** Triple staining of vGlut3, Tdtomato and Otoferlin in cochleae of control Plp1-Ai9 (A-A’’’) and Plp1-TAT (B-B’’’) mice at P42, both of which are treated TMX at P0 and P1 and TMP at P3 and P4. Arrows in (A-A’’’) and (B-B’’’) mark one IBC/IPh that is Tdtomato+/vGlut3-/Otoferlin-, and one new IHC that is Tdtomato+/vGlut3+/Otoferlin+, respectively. **(C-D’’)** Double staining of Myo7a and Tdtomato in cochleae of Plp1-Ai9 (C-C’’) and Plp1-TAT (D-D’’) mice. Likewise, arrows in (C-C’’’) and (D-D’’) mark one IBC/IPh that is Tdtomato+/Myo7a-, and one new IHC that is Tdtomato+/Myo7a+, respectively. **(E-F)** Quantification of new IHC numbers (E) or percentage (F) of new IHCs among all Tdtomato+ cells, using either Myo7a or vGlut3 as a marker at P42. Scale bar: 20 μm (B’’’ and D’’).

